# No supergene despite social polymorphism in the big-headed ant *Pheidole pallidula*

**DOI:** 10.1101/2022.12.06.519286

**Authors:** Emeline Favreau, Claude Lebas, Eckart Stolle, Anurag Priyam, Rodrigo Pracana, Serge Aron, Yannick Wurm

**Affiliations:** Organismal Biology Department, School of Biological and Chemical Sciences, Queen Mary University of London, Mile End Road, London, E1 4NS, United Kingdom; Association Roussillonnaise d’Entomologie, France; Center of Molecular Biodiversity Research, Zoological Research Museum Alexander Koenig, Leibniz Institute for the Analysis of Biodiversity Change (LIB), Adenauerallee 127, 53113 Bonn, Germany; Service Evolution Biologique & Ecologie, Université Libre de Bruxelles, Bruxelles, Belgium

## Abstract

Ant colonies ancestrally contained one queen and her non-reproductive workers. This is also the case for many but not all colonies of the Mediterranean big-headed ant *Pheidole pallidula*. Indeed, this species also has a derived form of social organization with multiple reproductive queens in the colony. The co-existence of two social forms also independently evolved in three other lineages of ants. In each of those lineages, variants of a supergene region of suppressed recombination determine social form. This is likely because supergene regions can link advantageous combinations of alleles from multiple loci. We thus hypothesized that a supergene region also determines colony queen number in the big-headed ant. To test this, we performed extensive population genetic analyses and genomic comparisons. We find no evidence of a supergene-like region with differentiation between single- and multiple-queen colonies. Our results show that a complex social polymorphism can evolve and be maintained without supergenes.

## Introduction

Many species include individuals with alternate discrete phenotypes that co-exist within the same population. This variation can be controlled genetically (polymorphism), or can occur in response to environmental conditions (polyphenism). The existence of complex polymorphisms and polyphenisms has long presented a conundrum for evolutionary biologists (Mayr 1963; Stearns 2010). Indeed, gene flow can cause alleles that benefit individuals with one phenotype to be carried by individuals of an alternative phenotype, where these alleles can be maladaptive. Several resolutions exist to this type of evolutionary antagonism. One that has been extensively discussed is the evolution of supergene regions of suppressed recombination (Stearns 2010; Thompson and Jiggins 2014; Gutiérrez-Valencia et al. 2021). Supergene regions emerge when the alleles contributing to a given phenotype become linked, for instance through the inversion of part of a chromosome. Supergenes allow linked alleles to be inherited as a unit and to be shielded from alleles that contribute to the alternate trait (Kirkpatrick and Barton 2006). High-throughput sequencing has enabled genome-wide comparisons between morphs in many species and thus led to the discovery of many examples of supergenes (*e.g*., S-locus in *Primula* heterostyly (Huu et al. 2016), P-locus in *Heliconius* Batesian mimicry (Joron et al. 2006; see review by Gutiérrez-Valencia et al. 2021). However, we still know little about how supergenes emerge, or whether they can arise from non-genetic polyphenisms.

Ants are excellent models for studying the evolution of complex phenotypes. They ancestrally have one queen per colony, yet many ant species have transitioned to having exclusively multiple-queen colonies (Hughes et al. 2008), and some species include both single- and multiple-queen colonies (Boulay et al. 2014). Differences between social forms are best-known in the red fire ant *Solenopsis invicta*. In this species, queens in single-queen colonies have larger abdomens, and workers are more aggressive than in multiple-queen colonies. Furthermore, young queens from single-queen colonies fly far from their original nest for mating, then independently found new colonies. In contrast, queens forming multiple-queen colonies rejoin their original nest after mating closeby, and a new multiple-queen colony forms when a group of queens and workers leave on foot. In *S. invicta* and other species, the multiple-queen social form is favored when dispersal risks are high and independent colony founding success is low (Hölldobler and Wilson 1990; Ross and Keller 1995).

Genetic comparisons between single- and multiple-queen colonies have been performed in three lineages of ants that are only distantly related, with most recent common ancestry between any pair of lineages being 119 million years ago (Blanchard and Moreau 2017). In all cases, social organization is controlled by supergene regions of suppressed recombination. The supergene regions evolved independently in each lineage and share no homology. The three supergene regions each include hundreds of genes; they span 8 megabases (Mb) in *Pogonomyrmex californicus* (Errbii et al. 2021), 9 Mb in *Formica selysi* (Purcell et al. 2014), and at least 20 Mb in *S. invicta* (Stolle et al. 2022). Sequence and gene expression differences between the two supergene variants of *S. invicta* (Pracana et al. 2017; Martinez-Ruiz et al. 2020; Stolle et al. 2022) are consistent with the theory that supergene architecture evolves to protect specific combinations of alleles from recombination (Kirkpatrick and Barton 2006; Thompson and Jiggins 2014; Rubenstein et al. 2019).

Finding that supergenes encode social polymorphism in each of the three ant lineages suggests that this genetic architecture is required for the evolution or maintenance of social polymorphism. However, the lack of homology between the supergene regions also suggests that multiple molecular pathways exist for encoding social polymorphism.

To understand whether the co-existence of single- and multiple-queen colonies requires a supergene region, we explored the genetic basis of variation in social organization in the Mediterranean big-headed ant *Pheidole pallidula*. This species is a distant relative to the other examined ant lineages, its most recent common ancestry being 51 million years ago with *Solenopsis* (Kumar et al. 2017). In the big-headed ant, single-queen and multiple-queen colonies co-exist in the same geographic areas (Aron et al. 1999). The two social forms differ in mating and dispersal strategies similarly to what occurs in *S. invicta* (Fournier et al. 2016). We generated population genomics data from the big-headed ant and used two types of analysis to test whether any segment of its genome shows the types of patterns that would be expected if a supergene controlled social form: we looked for signs of allelic differentiation and for signs of differences in genome coverage. We found population-specific differences in allele frequencies between social forms, but intriguingly no evidence of a supergene. Our results show that supergene architecture is not required for alternate social phenotypes, and highlights the types of allele frequency differences that could precede supergene evolution.

## Results

### Paucity of consistent genotype differences between single- and multiple-queen colonies

To determine whether a supergene contributes to social polymorphism in *P. pallidula*, we collected workers from 108 colonies across three populations: Bruniquel (Southern France), Vigliano (Central Italy) and Iberia (Spain) (Figure 1a, Supplementary Figure 1). We determined the social form of each colony by genotyping six polymorphic microsatellite loci on at least 8 workers per colony (Supplementary Table 2). Both social forms were present in each population, with 37 single-queen and 71 multiple-queen colonies overall (Supplementary Table 3). To enable genome-wide comparisons, we first constructed a long-read *de novo* genome assembly for this species (N50 length: 588 kb; total length: 287 Mb; BUSCO completeness: 98.8%; Supplementary Information, Supplementary Figure 2, Supplementary Tables 1 and 8). We subsequently sequenced the genome of one worker per colony using pairs of 150 bp reads (Supplementary Table 9), obtaining 1,300× genome coverage overall. From these data we identified 812,760 high-confidence Single Nucleotide Polymorphisms (SNPs; Supplementary Table 4).

**Figure 1:**
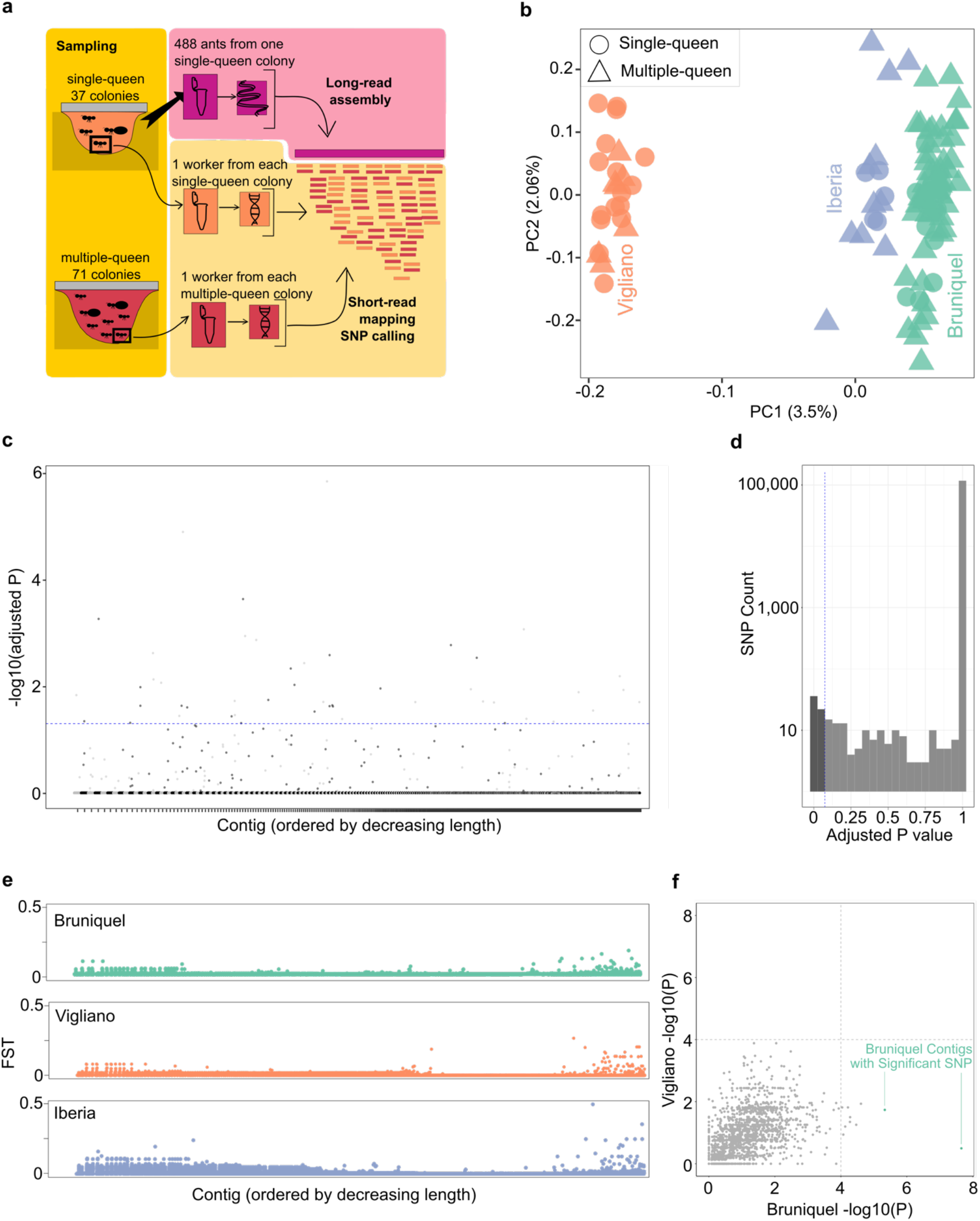
No evidence that a supergene region is genetically differentiated between single- and multiple-queen colonies. **a)** Experimental design: 108 sequenced workers, one per colony, originating from three populations containing both single-queen and multiple-queen colonies. **b)** The first two principal components based on 121,786 polymorphic SNPs separate populations but not social forms. **c)** Association tests across the whole dataset: out of 121,786 SNPs, 48 were significantly associated with social form (Fisher’s exact tests, Bonferroni adjusted P < 0.05, above the blue dashed line). SNP color alternates by contig; the 2,514 contigs are ordered by length. **d)** Distribution of Bonferroni-corrected *P* values for 121,786 Fisher’s exact tests of association with social form. The significant associations (below 0.05, dashed line) are represented in dark grey. **e)** F_ST_ between social forms within each population (sliding 30kb windows, or entire contig when smaller). The 2,514 contigs are ordered by length. There is no common region for which the three populations have a high F_ST_. High F_ST_ in the shortest contigs is a likely artifact of greater noise. f) The contigs with the strongest genetic differentiation between social forms in one population are not the most differentiated in the other population (Pearson’s correlation r = 0.443). 1,748 contigs containing SNPs within each population are represented by the most significant *P* value from each population (Fisher’s exact test on SNP data, raw *P* value). The two green contigs contain SNPs that are significantly associated with social organization in the Bruniquel population.

We performed four complementary lines of analyses of these SNPs to find evidence of allelic differentiation between individuals from single-queen and from multiple-queen colonies. If a supergene region were associated with social form, we would expect many closely-linked loci to be consistently differentiated between colony types and across the three populations. Our approach was guided by simulations of populations carrying supergene loci of *S. invicta* and of *F. selysi*. These simulations showed that our approach and the amount of genetic variation in our dataset should be sufficient to detect a potential supergene (Supplementary Information, Supplementary Figures 12, 13, 14 and 15).

We first performed principal component analysis of all retained SNPs. This shows that the first two principal components, which each explain less than 4% of the genetic variation in our dataset, separate samples by geographical region (Figure 1b). Similarly, none of the first 12 principal components show a clear association with social form (Supplementary Figure 5), suggesting that no large set of SNPs are systematically associated with social form.

We subsequently performed a genome-wide comparison between all individuals from single- and multiple-queen colonies. For this, we initially subset our dataset to the 121,786 SNPs that were polymorphic in all three populations and that had been genotyped in at least 75% of the samples. This subset includes a variant every 2,463 bp, a sufficient density to identify the smallest known supergenes, such as the stickleback fish supergene (200 kbp; Erickson et al. 2018) or the *Primula S* locus (300 kbp; Li et al. 2016). Forty-eight SNPs were significantly associated with social form (Fisher’s exact test *P_adj_* < 0.05, Bonferroni correction, Figure 1c). However, these SNPs did not show the characteristics expected if they were located within a supergene: the significant SNPs were scattered across the genome; no particular genomic regions were overrepresented (Supplementary Figure 6, Supplementary Tables 5 and 6). Analogous comparisons between single- and multiple-queen colonies using all variable SNPs or all Bruniquel SNPs yielded qualitatively similar results (Supplementary Information, Supplementary Figures 7, 8 and 9).

Third, we hypothesized that a potential supergene may be detectable only within one population because of supergene turnover or population-specific genetic variation, such as drift or recent local adaptation (Shi et al. 2020). To test whether we could detect population-specific evidence of a supergene, we performed an additional population-specific genome-wide comparison for each of our three populations. We found significant associations between SNPs and social form in the Bruniquel population only (20 SNPs, Fisher’s exact test *P_adj_* < 0.05, Bonferroni correction, Supplementary Figures 8 and 9). However, these SNPs were scattered across the genome, and only three of them were also variable in the other populations. Furthermore, the regions surrounding significant loci showed no trend for association with social form in the other datasets (Figure 1f, Supplementary Figure 10). Finally, we calculated the fixation index F_ST_ to test whether there is genetic differentiation between social forms within each population. Between *Solenopsis* siblings carrying alternate supergene variants, the average F_ST_ in the supergene region is 0.9, while between *Formica* colonies with alternate social forms, F_ST_ is in some species as high as 0.8 within the supergene region, and in any case at least 2-fold higher than the genome average (Pracana et al. 2017; Brelsford et al. 2020). We found that the genome-wide F_ST_ between social forms was consistently low within each *Pheidole* population (F_ST_ < 0.25 in each analysis with 30 kb window sizes), with only few isolated population-specific regions of F_ST_ peaks that deviated from the genome-wide distribution (Figure 1e). Thus potential genetic differences between social forms are weak and population-specific.

### Lack of coverage differences between single- and multiple-queen colonies

We hypothesized that if a supergene truly controls social form in *Pheidole*, the absence of signal in the previous analyses could be due to differences in the ability to map DNA sequence reads from the supergene to the reference genome. Two reasons could explain this. First, the supergene variant associated with multiple-queen colonies could be so divergent from the reference genome haplotype that read alignment and subsequent SNP identification becomes impossible with standard bioinformatic tools. Such divergence is analogous to the divergence between some sex chromosomes; indeed, some sex chromosomes can be identified as regions with low coverage in the heterogametic sex (Carey et al. 2022). Alternatively, social form could be controlled by a hemizygous supergene that is present in multiple-queen colonies but absent from single-queen colonies. This would make it comparable to the supergene controlling heterostyly in *Primula* primroses, with a hemizygous insertions present exclusively in one of the two morphs (Li et al. 2016). Creating a genome reference from each morph can enable the recovery of these types of supergenes (Vekemans et al. 2021).

We used three approaches to test whether the samples taken from multiple-queen colonies included sequences that are absent from the single-queen reference assembly (Supplementary Figure 11a). First, we produced *de novo* assemblies from reads that did not map to the single-queen reference assembly. The resulting contigs were all short and are likely to represent standard copy-number variation (Supplementary Table 7 and 10). Second, we identified contigs with extreme high coverage of reads either from single-queen colonies (top part of Figure 2) or from multiple-queen colonies (bottom part of Figure 2), to test whether the single-queen reference assembly may include duplicated segments such as pseudogenes associated with social form. We found that such contigs are short (median length of 1,986 bp) and carry none of the 48 SNPs significantly associated with social form in all populations. Third, we calculated for each sample the proportions of reads that map to the single-queen reference assembly, to determine if there may be loci present in many individuals that are absent from our reference. We found no systematic difference in mapped read proportions between social forms nor GC content (Supplementary Figures 3 and 4). In sum, none of these additional analyses suggested that a supergene region is associated with social form in this species.

**Figure 2:**
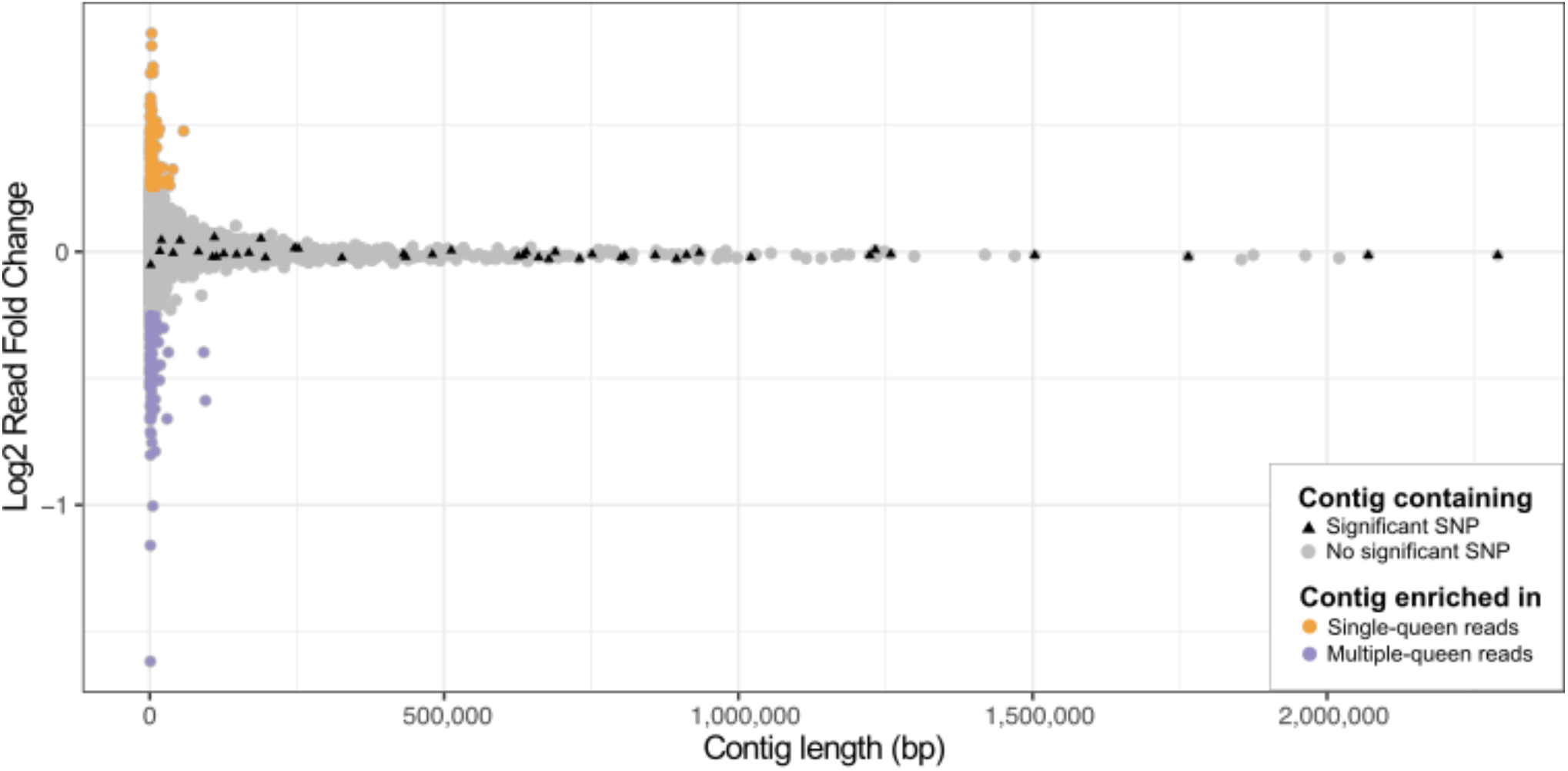
Relative sequencing coverage of reads from the two social forms for each contig of the reference genome. Contigs are ordered by length on the x axis. Contigs that are enriched in either single-queen colony reads (orange, top y axis) or multiple-queen colony reads (purple, bottom y axis) are very short (left x axis) and lack SNPs significantly associated with social form (triangles).

## Discussion

The Mediterranean big-headed ant *Pheidole pallidula* is a socially polymorphic species in which some colonies have the ancestral social form with a single queen, while other colonies have a derived social form with multiple queens. To test whether a supergene region determines social form in this species, we performed genome-wide comparisons across 108 colonies. We found some genetically differentiated loci between single-queen and multiple-queen colonies, but these loci are spread throughout the genome, with no evidence that they are linked together to form a supergene. These findings indicate that an alternate mechanism determines whether a colony of the big-headed ant will have one or multiple queens.

Our results suggest that social form in this species has an environmental and a genetic component. Ecological work has identified environmental conditions in which multiple-queen colonies are favored. In particular, intense competition over space and food resources can change population structure (Sundstrom 1995), and is associated with multiple-queen colonies (Herbers 1986; Boomsma et al. 2014), polydomal colonies (Burns et al. 2019), and supercoloniality (Helanterä 2022). Some species may have more plasticity in social form than others in responding to competition. Our finding alleles associated with social form highlights which types of changes may facilitate this plasticity, for example by increasing the likelihood of a newly mated queen to join an established colony, or lowering worker aggression.

We found species-specific and population-specific associations between genotype and social form, which indicates that social form may have a genetic component in *P. pallidula*. Some of these loci may have stood out in our analysis due to genetic differentiation that is neutral, linked to demography, or otherwise unrelated to social form.

Nevertheless, the existence of alleles associated with social form in this species has an important implication for understanding the early steps in supergene evolution. Alleles at some of these loci could increase the likelihood that a colony accepts multiple queens, while alleles at other loci could increase survival of multiple-queen colonies. In this situation, if allelic effects are strong enough, selection could favor locking the best allelic combinations together. This could, for example, occur through a chromosomal inversion event and would represent a hallmark in supergene evolution (Kirkpatrick and Barton 2006; Thompson and Jiggins 2014). Further studies of other *Pheidole* populations or species may thus show patterns that differ from ours.

Overall, our study suggests that the evolution and maintenance of social polymorphism can occur without a supergene. Further work, ideally combining long-term field observation, cross-breeding experiments and multi-generational analyses, will produce a more extensive survey of the potential mechanisms at play in *P. pallidula*, and, ultimately, of the evolution of social organization in the ants.

## Data availability

Raw reads and analysed data for both the short reads and long reads are available at NCBI PRJNA721572 and PRJNA729742. All other data supporting the findings of this study are available within the paper and its supplementary files (Supplementary Tables, Supplementary Information), and at https://wurmlab.com/data. All scripts are available on Github at https://github.com/EmelineFavreau/Pheidole_pallidula_social_polymorphism.

## Acknowledgements

We thank Xavier Espalader, Nilo Ortiz de Zugasti Carron, Pedro Lorite Martínez, Janine Remoué and Carlos Martinez Ruiz for assisting in the sampling effort; Richard A. Nichols, Christophe Eizaguirre and Max Reuter for advice and discussion. This work was supported by the Natural Environment Research Council (grant NE/L002485/1 to E.F. and NE/L00626X/1 to Y.W.) and the Biotechnology and Biological Sciences Research Council (grant BB/K004204/1 to Y.W.). This research utilised Queen Mary’s Apocrita HPC facility, supported by QMUL Research-IT.

## Author contributions

E.F. contributed to the study design, data collection, laboratory experiments, data analysis, manuscript writing; C.L. contributed to data collection; E.S. contributed to laboratory experiments, manuscript writing; A.P. contributed to data analysis; R.P. contributed to the study design, manuscript writing; S.A. contributed to data collection; Y.W. contributed to the study design, data collection, manuscript writing.

