## Supplementary Information for "No supergene despite social polymorphism in the big-headed ant *Pheidole pallidula*"

### Wet Lab Protocols

*Protocol 1: Microsatellite sequencing preparation*

**Ant preparation for 24 ants (6 colonies):**

1. Take ant samples out of freezer
2. Turn on the hot plate (55 degrees)
3. Label screwtop tubes (n=24, cap and side)
4. Add 1g beads in each tube
5. Add 1 ant in each tube
6. Dry ethanol on hot plate (55 degrees)
7. Repeat these steps for each ant
8. Put back the ant samples in the freezer

**DNA extraction protocol (Gadau, 2009) for 24 ants:**

1. Make 12.5 mL of Chelex
    a. Collect Chelex (Chelex 100 (Biorad)), deionised water, bleach, magnetic stirrer, spatula, 1 15 mL falcon tube
    b. Bleach magnetic stirrer and spatula
    c. Put 12.5 mL of deionised water in falcon tube
    d. Add 0.63 g of Chelex with spatula into falcon tube
    e. Add the magnetic stirrer in falcon tube, mix for 1 min

2. Add 360uL Chelex to each sample tube with 1000 µl tip pipette. Make sure to take same amount of Chelex beads each time (vortex)
3. Turn on the hot plate to 57 °C. Take proteinase K out of freezer.

4. Shake samples at full power on FastPrep homogeniser for 60 s then 60 s rest then again 60 s shake
5. Spin tubes for 1 minute at full speed (centrifuge)
6. Add 1 µl proteinase K to each tube
7. Put the tubes on 57 °C hot plate for 1 hour. Turn on another hot plate on 95 °C.
8. Put the tubes on 95 °C hot plate for 5 min
9. Spin tubes for 15 min at full speed (centrifuge)
10. Meanwhile label set of Eppendorf tubes.
11. Transfer supernatant into new tubes (rough volume: 200 µl)
12. Store extraction in -20 °C freezer

**PCR for one plate**

1. Make primer mix of 6 primer pairs (following Qiagen protocol in Type-it Microsatellite PCR kit: forward labelled primers from 10× conc stock (50 pmol/µL), reverse primers from 10× conc stock (50 pmol/µL), TE buffer)
2. Prepare PCR plate notes (which sample in which well?)
3. Program the PCR machines following the Qiagen kit protocol
    a. 5 min at 95 °C
    b. 30 s at 95 °C
    c. 90 s at 61 °C (temperature specific to *pallidula*)
    d. 30 s at 72 °C
    e. Number of cycles (35 here, because small amount of DNA template)
    f. 30 min at 60 °C
4. Collect Type-it mastermix, DNA, RNase-free water, primer mix, DNA samples, 1 eppendorf, PCR plate and seal. Pipette all up and down.
5. Prepare reaction mix in eppendorf
    a. Primer mix: 122.18 µl
    b. Type-it mastermix: 706 µl
    c. Water: 439.64 µl
6. In PCR plates, add 11 µL reaction mix in each well.
7. In PCR plates, add 1 µl DNA. Mix well.
8. Apply seal on plate (secure properly).
9. Run the PCR.
10. Store PCR products in -20 °C freezer

*Protocol 2: Phenol-chloroform protocol for short read library preparati***on**

*Removing the ethanol from ant sample, preparing solution*

1. Leave the whole ant in an eppendorf tube with 1.5 mL deionised water for an hour on the bench at RT (room temperature).
2. Meanwhile, make up a sodium acetate-ethanol solution. In a falcon tube, add 40 mL 100% ethanol and 2 mL 3M sodium acetate solution. Keep it on the bench for later.

*Breaking the cuticle, digesting proteins*

3. Set the heat block at 55 °C.
4. In an eppendorf, add 350 μl CTAB, 10 μl Proteinase K (10 mg/mL). Transfer the ant from the water eppendorf to a clean blue tissue to remove as much water as possible, then put it in the CTAB + Prot K eppendorf. Use a pestle to reduce the ant into small fragments - no full appendage should be identified. Incubate the ant solution on the heat block for an hour.

*Taking the proteins out of the solution, keeping the DNA in the water solution*

5. Place phenol, phenol:chloroform, 1000 μl pipette tips and centrifuge in the fume hood. Keep gloves at all times when handling equipment and samples in the fume hood. As soon as your hands are out of the fume hood, change gloves.
6. In each eppendorf with the ant solution, pipette 350 μl of the lower phase of phenol. Close the lid, shake lightly 10 times. Load the samples in the centrifuge and leave to rest 10 minutes before activating the centrifuge (10 minutes, full power).
7. Meanwhile, label 2 series of eppendorf tubes and add 350 μl of chloroform in each. Label a 3rd series of eppendorf tubes and add 1 mL sodium acetate-ethanol solution (prepared earlier). Put series 1 and 2 on the bench near centrifuge, and series 3 in the freezer.
8. When the centrifuge is finished, take each sample and pipette the upper phase into series 1 of eppendorf prepared earlier (with 350 μl of chloroform). Close the lid, shake lightly 10 times. Load the samples in the centrifuge and leave to rest 5 minutes before activating the centrifuge (10 minutes, full power).
9. When the centrifuge is finished, take each sample and pipette the upper phase into series 2 of eppendorf prepared earlier (with 350 μl of chloroform). Close the lid, shake lightly 10 times. Load the samples in the centrifuge and run it without delay (5 minutes, full power).
10. When the centrifuge is finished, take each sample and pipette the supernatant into series 3 of eppendorf prepared earlier (with 1 mL of sodium acetate - ethanol). Expect to see a DNA pellet. Store in -20 °C freezer for an overnight precipitation.
11. Clean equipment that was in the fume hood: soapy water for tube tray, bleach on pipette. Put centrifuge back on the bench.

*After overnight precipitation, 2x ethanol wash*

12. Centrifuge at full speed for 20 min, putting the tubes with hinges outwards. A DNA pellet should be visible.

13. Remove as much supernatant as possible: turn the tube on a waste falcon tube, watching out the pellet (don't let it drop). Wipe opening on blue paper.
14. Add 500 µl of 70% ethanol. Close lid, swing twice.
15. Centrifuge at full speed for 20 min, putting the tubes with hinges outwards.
16. Remove as much supernatant as possible: turn the tube on a waste falcon tube, watching out the pellet (don't let it drop). Wipe opening on blue paper.
17. Add 500 µl of 70% ethanol. Close lid, swing twice.
18. Centrifuge at full speed for 20 min, putting the tubes with hinges outwards.
19. Remove all supernatant: turn the tube on a waste falcon tube, watching out the pellet (don't let it drop). Wipe opening on blue paper. Air dry for 10 minutes.
20. Once no ethanol remains in the tube, dissolve the DNA pellet in 20 - 30 µl of low TE buffer or deionised water.

*Protocol 3: DNA cleaning step before Illumina library preparation*

This protocol was used right after the phenol-chloroform extraction. Modifications in the extraction protocol: 1 g of ceramic beads in screw cap tubes on Fastprep (no ant part should be identifiable after homogenization), only one ethanol wash (ethanol on ice), air dry, molecular water for eluting.
Then we use part of the GenElute kit for cleaning, by starting at step 4 of the official protocol.

**Material**

- DNA extraction (volume = 20 µl in molecular water)
- Molecular water
- 100 % ethanol
- GenElute™ Mammalian Genomic DNA Miniprep Kit from Sigma Aldrich (G1N70)

**Preparing steps:**

1. Prepare GenElute tubes (per sample: 1 pre-assembled GenElute Miniprep Binding Column, 3 2 mL collection tubes)
2. Prepare ethanol (per sample: 200 µl)
3. Prepare GenElute solutions if necessary

**Cleaning steps:**

1. Add 400 µl molecular water to the extraction.
2. *Column preparation*: Add 500 µL of the Column Preparation Solution to each pre-assembled GenElute Miniprep Binding Column (with a red o-ring) and centrifuge at 12,000× g for 1 minute. Discard flowthrough liquid.
3. *Prepare for binding*: Add 200 µL of ethanol (95–100%) to the lysate; mix thoroughly by vortexing 5–10 seconds. A homogeneous solution is essential.
4. *Load lysate*: Transfer the entire contents of the DNA extraction tube into the treated binding column from step 2. Use a wide bore pipette tip to reduce shearing the DNA when transferring contents into the binding column. Centrifuge at ≥6500 × g for 1 minute. Discard the collection tube containing the flowthrough liquid and place the binding column in a new 2 mL collection tube.
5. *First wash*: Add 500 µL of Wash Solution to the binding column and centrifuge for 1 minute at ≥6,500× g. Discard the collection tube containing the flow-through liquid and place the binding column in a new 2 mL collection tube.
6. *Second wash*: Add another 500 µL of Wash Solution to the binding column; centrifuge for 3 minutes at maximum speed (12,000-16,000× g) to dry the binding column. The binding column must be free of ethanol before eluting the DNA. Centrifuge the column for one additional minute at maximum speed if residual ethanol is seen. You may empty and re-use the collection tube if you need this additional centrifugation step. Finally, discard the collection tube containing the flowthrough liquid and place the binding column in a new 2 mL collection tube.

7. *First elution of DNA*: Pipette 150 µL of molecular water directly into the center of the binding column; incubate for 10 minutes at room temperature; centrifuge for 1 minute at ≥6,500× g to elute the DNA.
8. *Second elution of DNA*: Pipette 150 µL of molecular water directly into the center of the binding column; incubate for 10 minutes at room temperature; centrifuge for 1 minute at ≥6,500× g to elute the DNA. Current volume: 300 µL.
9. *Reduce volume*: Speed vac (4 °C) for a final volume of 30 µl.
10. Store in fridge in GenElute tube if WGS library prep will happen very soon. If not, I use low binding tubes.

### Supplementary Information

#### Sample collection

From each of 108 *Pheidole pallidula* colonies, 58 colonies were sampled in 2002-04 from France (Fournier et al. 2002) and Spain, and 57 were sampled in 2016-17 from France, Spain and Italy (Supplementary Table 3). Based on their geographic distribution, we expect all samples to be from the subspecies *Pheidole pallidula* (Seifert 2016). This was subsequently confirmed using mitochondrial barcoding (see below). We collected minor and major workers, either selected from within the colony, or attracted with bait outside the colony. All were stored in 100% molecular grade ethanol. We named three populations based on the location of the majority of samples: Bruniquel, Vigliano, Iberia (Supplementary Figure 1).

#### Microsatellite genotyping

We used microsatellite genotyping of workers from each colony to determine its social form. We first extracted DNA from eight workers of each colony, using a protocol based on a co-polymer solution (Gadau 2009). Briefly, we reduced each individual into small fragments on a FastPrep homogeniser (MP Biomedicals) for two two-minute cycles of 8,000 rpm separated by one minute of rest, in a 5% Chelex solution with 1 g ceramic beads. We then proceeded to the incubation and centrifuge steps as per the protocol. We evaluated DNA yields using fluorometry (Qubit 2.0, Life Technologies).

We then amplified six microsatellite loci, using species-specific markers (Fournier et al. 2002), with fluorescent forward primers (VIC *Ppal01T,* NED *Ppal33,* PET *Ppal84*, FAM *Ppal03*, FAM *Ppal73,* VIC *Ppal12*) and non-fluorescent reverse primers. We performed multiplex PCRs using Type-It PCR kit (Qiagen) with 1 µL of extracted DNA, following the manufacturer's cycling conditions modifying the annealing step (90 s at 61 °C) and the total number of cycles (35). Genotyping was performed on a 3730 DNA Analyser (Applied Biosystems); we subsequently called microsatellite alleles with GeneMarker (v.2.4.0, SoftGenetics). More details are provided in the Wet Lab Protocols section of this document.

For each colony, we estimated the number of queens by counting the number of alleles present at each microsatellite locus, with an average allele number per locus of 27.5 (Supplementary Table 2). With the assumptions that this species is singly-mated (Fournier et al. 2002) and that eight workers are a fair representation of the colony genotype, we use a simple rule: if more than three alleles are present at a locus within a colony dataset, or if the colony clearly contains workers from different fathers, then the colony has multiple queens.

We subsequently evaluated the level of potential sample outliers with a principal component analysis, inputting the genotypes of each colony in the R package adegenet (Jombart 2008; R Core Team 2014).

#### DNA extraction for Illumina library preparation and sequencing

We first extracted high-quality DNA from one representative worker from 39 single-queen colonies and from 76 multiple-queen colonies (Supplementary Table 3). We followed a phenol-chloroform extraction protocol (Hunt and Page 1995), with slight modifications: 10 µL Proteinase K were added to the CTAB step, and we omitted the NaCl-Tris-Cl step. We further cleaned the extractions using part of Sigma Aldrich GenElute™ Mammalian Genomic DNA Miniprep Kit protocol (from step 4; catalogue number G1N70) and reduced the extraction volume to 20 µL with an Evaporator centrifuge (Uniequip, Univapo 100H). More details are in the Wet Lab Protocols section of this document.

We then prepared individual libraries for whole-genome sequencing, using NEBNext® Ultra™ II FS DNA Library Prep Kit for Illumina (catalogue number E7805) and a combination of two primer sets (NEBNext® Multiplex Oligos for Illumina® Dual Index Primers Set 1 catalogue number E7600 and Set 2 catalogue number E7780). We performed the protocol in half volumes, to improve performance while reducing the costs (Tan and Mikheyev 2016). Each resulting library was controlled for fragment size (300 bp) using TapeStation High-Sensitivity tape (Agilent). 115 libraries were pooled in equimolar quantities. The final 15 nM pool was sequenced on three lanes of Illumina HiSeq 4000 with 150 bp paired-end reads (Genewiz). We obtained 2,762,930,432 short-read raw sequences (Supplementary Table 9, found on NCBI under PRJNA721572), which is the equivalent of more than 1,300× genome coverage (based on genome size estimate from Tsutsui et al. 2008). Each sample contributes to an average of 16× coverage, with one sample, E15, sequenced at 66× coverage.

##### Species identification

We performed species identification and all subsequent tasks on the QMUL Apocrita computing facility (King et al.). We downloaded every COI barcode sequence for the taxon "Pheidole" in the BOLD database (http://v3.boldsystems.org/), as well as the unique COI sequence of *Pheidole pallidula* from NCBI (GenBank: EF518381.1), whose sample originated from France (pers. comms Corrie Moreau). We reduced the number of BOLD sequences by collapsing redundant sequences (cd-hit-est v4.6.8, overlap c = 0.97, word size n=10, length of description in .clstr file d= 0). The final database file contained 600 BOLD sequences and one NCBI sequence. For each sample, we compared the Illumina raw reads (forward and reverse) against that database (Magic-BLAST v1.4.0 -dbtype nucl -parse_seqids (Boratyn et al. 2019)). To identify each sample, we assign the taxon name of the database sequences that bore the following criteria: 100% identity and the highest alignment score. All samples were identified as *P. pallidula*, with predictable regional proximity: our Italian and Corsican samples match BOLD barcodes from an Italian sample, French and Spanish samples match the NCBI barcode from the French sample.

##### Long read library preparation and sequencing

Prior to this study, no reference genome existed for this species and the most closely related genomes were from *Solenopsis invicta* (Wurm et al. 2011), *Atta cephalotes* (Suen et al. 2011) and *Acrymyrmex echinatior* (Nygaard et al. 2011), species that shared ancestry more than 51 million years ago Ward et al. 2015). To generate a reference genome sequence for *Pheidole pallidula*, we sequenced long reads with Oxford Nanopore (ONT) MinION from material extracted from males and workers of two single-queen colonies.

To extract high-molecular weight DNA from each sample, we first reduced the samples in pellets using either a hand pestle, or a tissueRuptor. We then applied the phenol-chloroform extraction method mentioned above, with special modifications to ensure the preservation of long molecules. We prepared six libraries based on two chemistry kits (2D and 1D$^2$); following ONT MinION protocols with several modifications (Supplementary Figure 2). We extracted the DNA using a phenol:chloroform method (Hunt and Page 1995). Briefly, after 1 h of wash in deionised water, each sample was ground with a pestle and digested in 350 µL CTAB buffer and 10 µL proteinase K enzyme. We then ran the solution through 350 µL of phenol:chloroform, followed by centrifugation and 2 subsequent chloroform washes. We transferred the final supernatant into 80% ethanol and 3M sodium acetate for an overnight precipitation step at −20 °C. The DNA precipitation was then washed twice with 70% ethanol and finally eluted in 20-30 µL of EB buffer. We quantified the DNA with fluorometry. We then followed a ONT protocol for library preparation, loaded the MinION sequencer and sequenced for 48 hours. In a research and development assay, we prepared each library with slightly different protocols, as summarised in Supplementary Figure 2. The libraries were subsequently sequenced following the manufacturer's instructions. Six sets of reads were obtained from the MinION runs (17×), basecalled with ONT Albacore version 2.1.7. We obtained raw reads with an average sequence length of 2.6 kbp (Supplementary Table 8) and an overall genome coverage of 17× (Supplementary Figure 2).

##### *De novo* assembly Ppal_gnE

We assembled all MinION reads, using Flye v2.4 (Kolmogorov et al. 2019) and a genome size estimation of 300 Mbp (Tsutsui et al. 2008). We used Pilon v1.22 (Walker et al. 2014) to "polish" the assembly, by reducing error rate and increasing contiguity. Pilon parameters were: --fix snps, indels --diploid. For this we used ten Illumina samples (I13-M, I02-M, I11-M, I23-M, I03-M, I12-M, I10-M, I18-M, I24-M, I14-M) that had been subsampled and cleaned. For the subsampling of 9.3× coverage per sample, we used seqtk (version 1.2, https://github.com/lh3/seqtk) for a combined coverage of 93×. For read cleaning, we first filtered the reads prior to use. First, we removed duplicates using clumpify.sh (from BBMap package, version 37; biostars.org). This removes duplicated reads regardless of origin (optical, PCR, other), which improves the efficiency of the polishing step. Second, we compared the mean base quality per cycle, per tile to the mean base quality of that cycle across all tiles to test for air-bubbles becoming trapped in the flowcell (sequencing.qcfail.com). For this, we obtained the difference between per-cycle mean base quality for a tile and the per-cycle mean base quality for all tiles from fastqc's (version 0.11.5; bioinformatics.babraham.ac.uk) text output. Where this difference was greater than 4, we changed the corresponding base in the reads to 'N'. This was done by creating a BED file of positions from the tile and cycle information and then using seqtk (version 1.2; https://github.com/lh3/seqtk) to convert bases at the positions specified in the file. Next, we considered that reads with multiple occurrences of low-quality bases may be problematic (Edgar and Flyvbjerg 2015). To eliminate such reads, we turned bases with quality score lower than 12 to 'N' using seqtk (reads with excessive Ns are removed in the next step). Third, we used bbmerge to correct overlapping read pairs. Fourth, we removed reads with mean quality threshold lower than 15 using htqc v0.90.8 (Yang et al. 2013). Fifth, we masked singleton kmers. Finally, we used cutadapt v1.13 (Martin 2011) to trim adapter sequences, to trim low quality bases from both 3' and 5' ends (thresholds were chosen specifically for each lane based on the sequence quality report generated by fastqc; see supplementary code repository), to trim any leading and trailing 'N's, and, after trimming, to eliminate reads shorter than 50 bp and those with more than 4 'N's.

We scaffolded Ppal_gnE using AGOUTI (Zhang et al. 2016) using publicly available RNAseq reads (GenBank: EF518381.1). We first mapped these RNAseq reads to each other with bwa (Li and Durbin 2009), and we generated an annotation file with MAKER (Cantarel et al. 2008) for AGOUTI input. We performed AGOUTI scaffolding with a minimum of five supporting RNA reads, with 318 contigs being scaffolded. This scaffolding step improved increased the contiguity of the genome from an N50 of 446,424 bp to 587,760 bp, measured by QUAST v4.6.1 (Gurevich et al. 2013)). This assembly N50 ranked among the top Hymenoptera genomes at the time of writing (Supplementary Table 1). It decreased the number of scaffolds from 4,130 to 3,954, a greater fragmentation than the expected 10 chromosomes in *Pheidole* (Lorite and Palomeque 2010). The length of this scaffolded assembly was 287 Mb, reasonably close to the flow cytometry-based estimate for the genome size of another *Pheidole* species (326 Mb, (Tsutsui et al. 2008)). We used BUSCO v3.0 (Simão et al. 2015) to estimate the gene completeness of the scaffolded assembly, which showed that 98.8% of 1,658 expected single-copy insect genes are present and complete, while only 0.7% are duplicated.

#### Reference-based analyses: mapping, variant calling, filtering, PCA, Fisher's exact tests

We performed a reference-based variant calling using the assembly Ppal_gnE and Illumina raw reads. We first mapped raw reads of each sample to the assembly using Bowtie2 v2.3.4 (Langmead et al. 2009), obtaining 108 BAM files with alignments private to each sample. Mapping quality was high for all samples retained for the analysis (Supplementary Figure 3) and GC content was consistent across samples (Supplementary Figure 4).

We then used FreeBayes v1.2.0 (Garrison and Marth 2012) with `--use-best-n-alleles 2` to call variants between the samples, obtaining 587,048 variants. We sorted and indexed the VCF file with BCFtools v1.8 (Li et al. 2009) and Tabix v0.2.5 (Heng Li 2011). We used VCFtools v0.1.15 (Danecek et al. 2011) to keep biallelic SNPs, with a minimum quality phred of 30 and minimum sample support of 75% (--remove-indels --minQ 30 --min-alleles 2 --max-alleles 2).

We initially investigated this variant dataset by conducting a PCA using PLINK v1.90b4.6 (Chang et al. 2015) (--allow-extra-chr --allow-no-sex --pheno --cluster) and visualised the results using R. The first ten principle components showed no clear clustering of samples by social form (Supplementary Figure 5).

We tested for association with social organization with a logistic regression in PLINK, first with town location coded in covariates (Supplementary Table 3). We also ran a logistic regression with PC1 and PC2 loadings as covariates that are proxy to geographical origins (Figure 1b in the main text). Using R, we corrected each resulting *P* value for multiple comparisons with Bonferroni correction.

We further filtered the variant database with VCFtools by keeping SNPs for which the minor allele count is greater than 1 in each population. This is because Fisher's exact test does not allow population structure as covariate. Using R, we corrected each resulting *P* value for multiple comparisons with Bonferroni correction. To investigate the association strength of the 48 loci that are significantly associated with social form in the main dataset, we assessed the frequency of the most common allele at each significant locus, in comparison with random loci found in the same contig (Supplementary Figure 6), inspired by *S. invicta* diagnostic enzyme frequencies between single-queen and multiple-queen colonies (Ross and Keller 1995).

We replicated the association tests for coding regions only (Supplementary Figure 7) and for each population (97,684 within-Bruniquel SNPs called between 69 samples, 48,613 within-Vigliano SNPs called between 23 samples, 71,052 within-Iberia SNPs). After Bonferroni correction, the Bruniquel population is the only population with SNPs that are significantly associated with social type (20 SNPs with $P_{adj} < 0.05$, Supplementary Figures 3 and 4). The degree of association of these SNPs is correlated between Bruniquel and Vigliano (Pearson's Correlation on unadjusted *P* values, r = 0.44, t = 20.651, df = 1746, p-value < 2.2e-16, Supplementary Figure 10). We also calculated FST values between social forms for each population using R package PopGenome v.2.7.5 (Pfeifer et al. 2014).

#### Simulations of association test with supergene region

To test the robustness of our analysis design, we performed several simulations using as input 3,757 real *Pheidole* SNPs (100% sample-coverage, in coding regions, within-population polymorphic, from 108 diploid workers).

We first performed a unit test with a simulation in which one simulated variant (homozygote in single-queen samples, heterozygote in multiple-queen samples) was added to the real SNPs. We ran a Fisher's exact test (allele count in two sample categories: single-queen and multiple-queen) using PLINK, adjusted for multiple comparisons in R (p.adjust, Benjamini & Hochberg method (Benjamini and Hochberg 1995)). This simulated SNP was by far the most significant (Supplementary Figure 12).

Then, to test whether our analysis would detect a realistic supergene, we performed a scaled simulation that replicates the model of *S. invicta*. Based on previous work (Pracana et al. 2017), 2.5% of all SNPs are fixed in *S. invicta* supergene region. We therefore added 95 simulated SNPs to the dataset, represent what we expect from the supergene system: homozygous SNPs for all single-queen and 1/3 of multiple-queen samples, heterozygous SNPs for 2/3 of multiple-queen samples (Buechel et al. 2014). We then ran Fisher's exact tests and adjusted for multiple comparisons. All simulated SNPs are associated with social form ($P_{adj}$ value < 0.05), strongly segregated away from the real dataset points (Supplementary Figure 13).

Finally, we simulated a dataset that replicates the model of *F. selysi*. Based on previous work (Purcell et al. 2014), 3.7% of all SNPs are fixed in the supergene region. We therefore added 139 simulated SNPs to the dataset, represent what we expect from the supergene system: homozygous SNPs for all single-queen, 68% heterozygote multiple-queen samples and 32% multiple-queen samples that are homozygous for the alternative allele (Supplementary Figure 14). All simulated SNPs are associated with social form ($P_{adj}$ < 0.05), strongly segregated away from the real dataset points. As predicted, the Fisher's exact test detects the *Formica* SNPs (in which there is strictly no homozygote for reference in multiple-queen samples) more easily than the *Solenopsis* SNPs, with respectively *P* values of 7.7E-9 and 6.4E-18.

Additionally, we investigated the effect of misgenotyping some of the samples on our analysis. We set the alternative social type to 10% of our samples and investigated the subsequent association tests. We find that a large proportion of the 48 real SNPs are always recovered as significant (i.e. true positive) in the simulations (Supplementary Figure 15).

##### Testing for differences in coverage between social forms

Some supergenes take the form of duplicated or a deletion of a genome segment. We thus tested whether there were differences in sequencing read depth between individuals from the two colony types (Supplementary Figure 11a). For each Bruniquel sample, we measured the alignment read depth using BEDtools genomecov (Quinlan and Hall 2010) and calculated average $\log_2$-fold changes between social forms. No entire genomic contigs showed significant differences in coverage between the two social forms (Figure 2; Supplementary Figure 11b). An analogous analysis of 5,000 nucleotide non-overlapping windows that include coding sequence and genetic variation revealed that 36 (1.6 %) of 2,296 such windows showed a significant difference of read coverage between social forms (Kolmogorov-Smirnov tests, $P_{adj}$ < 0.05; Benjamini-Hochberg adjustment; Supplementary Figure 11c). There was no general trend in direction with 17 of these windows having greater coverage in individuals from single-queen colonies, and 19 in multiple-queen colonies. None of these windows were on the same contig, indicating that they do not together form a supergene of 10,000 nucleotides or longer. We suspect that these small differences spread throughout the genome represent typical standing copy-number variation that may be segregating due to genetic drift. In conclusion, we find no large segment of the genome with read coverage differences between social forms.

##### Assembly of multiple-queen reads that do not map to the single-queen reference

We searched for genomic sequences representing a supergene present in multiple-queen samples but not in single-queen samples, and therefore absent from the reference genome assembly. Our null hypothesis is that any region that is unique to multiple-queen samples is not biologically relevant to social organisation, instead representing either non-assembled repeats or biological contaminants with little sequence similarity with assemblies from other ant species. To obtain genomic regions that are unique to and common to all multiple-queen samples, we first obtained reads from multiple-queen samples that did not map to the single-queen reference genome Ppal_gnE, using Samtools. We assembled these reads (unique to multiple-queen samples) into scaffolds using Spades version 3.12.0 (Nurk et al. 2013), the assembler that is tailored for small genomes. Using Bowtie2, we mapped the reads unique to multiple-queen samples to these scaffolds, then filtering with bedops and bedtools for regions that are common to all 73 multiple-queen samples. We blasted the 161 regions longer than 1,000 bp against Hymenoptera (NCBI blastn) and against *Solenopsis invicta*. Most of the regions (150 out of 161) match parts of sequences (average query cover: 28 %) from ants but none match a *Solenopsis* supergene region (Supplementary Table 10). It is thus likely that they represent frequently observed types of structural polymorphisms: copy number variation and/or repetitive elements. In conclusion, we do not find evidence of a supergene in the unique regions of multiple-queen assembly.

### Supplementary Figures

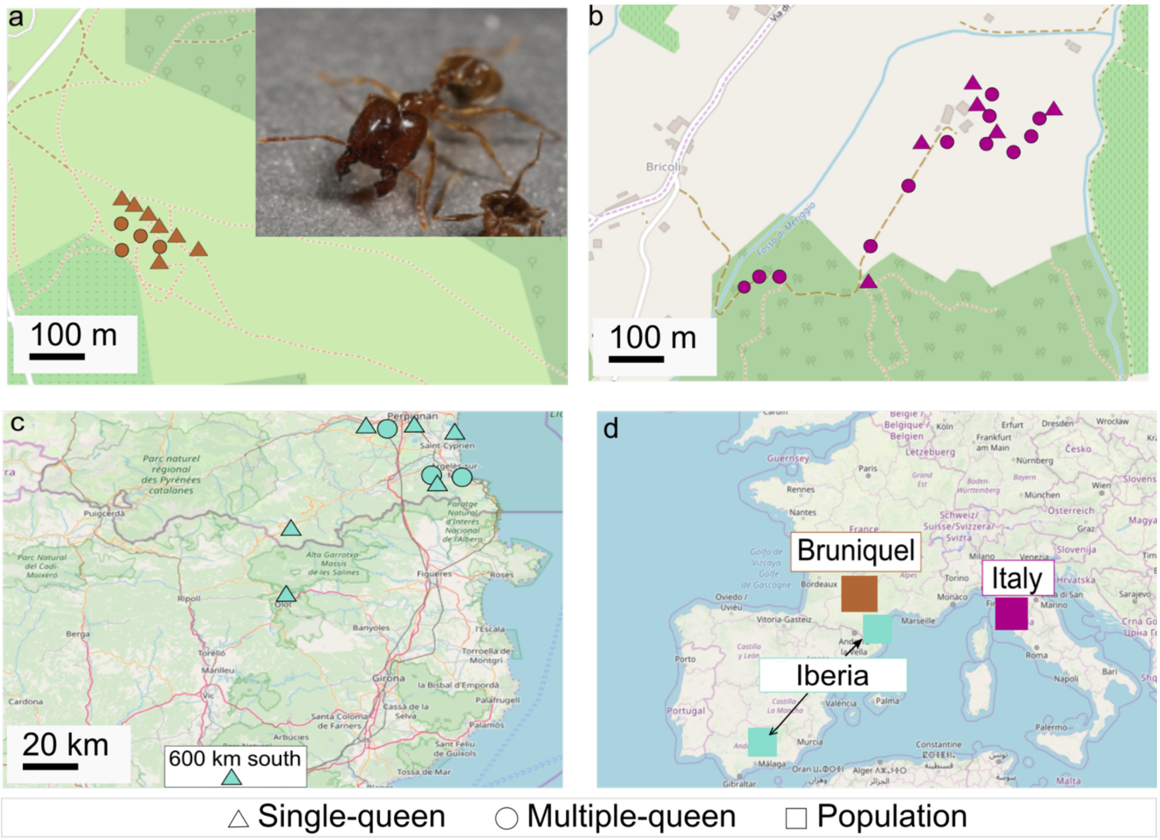

**Supplementary Figure 1: Geographical location map of samples.**
**a)** Bruniquel population: 53 polygynous and 16 monogynous samples. Insert: *Pheidole pallidula*'s big-headed soldier
**b)** Italy population (Vigliano): 7 polygynous and 16 monogynous samples.
**c)** Iberia population: 11 polygynous and 5 monogynous samples.
**d**) Overview of the populations.
Background map (OpenStreetMap contributors)

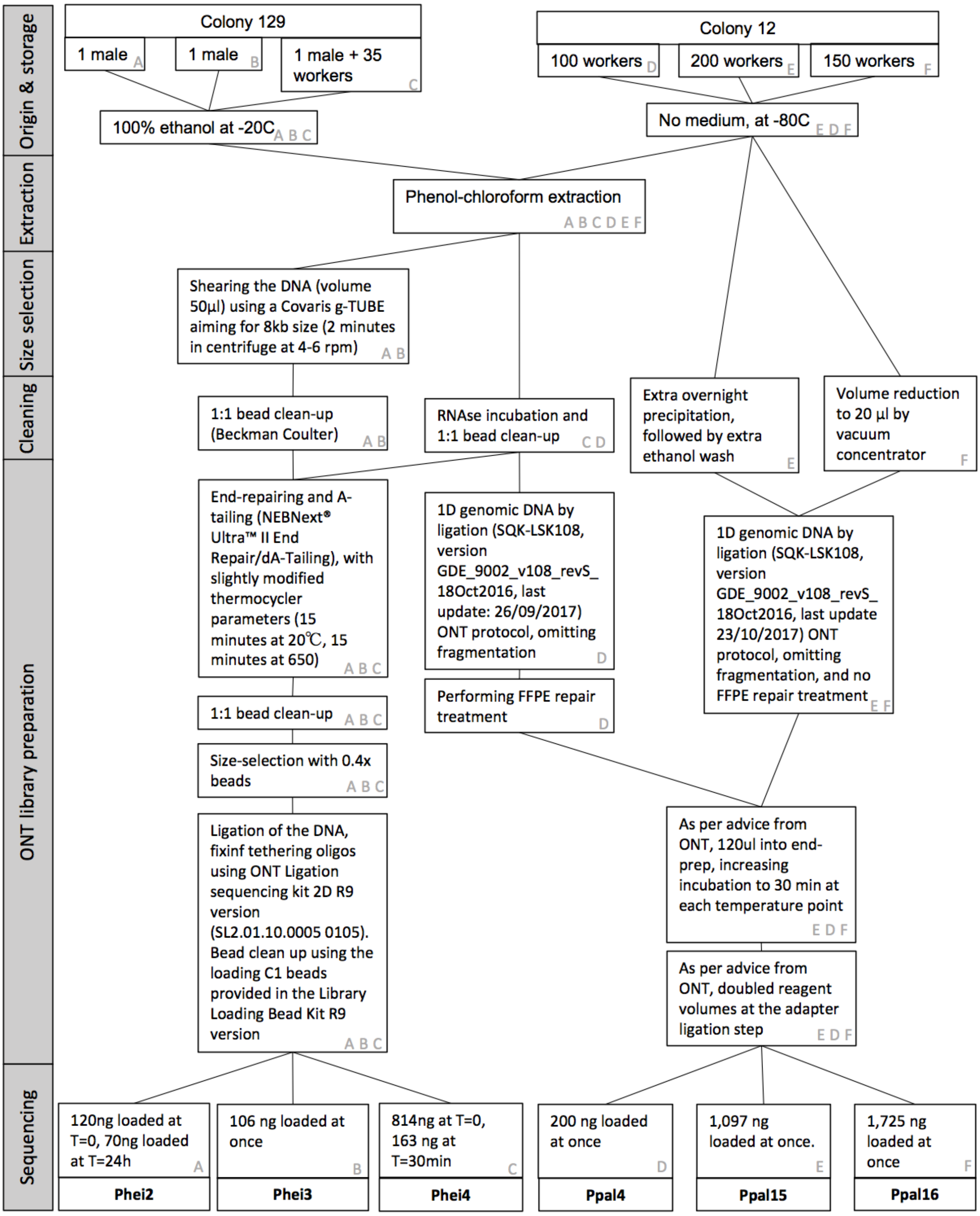

429 **Supplementary Figure 2: MinION protocols.**

430 Each flowcell received a unique combination of protocols (labelled A to F).

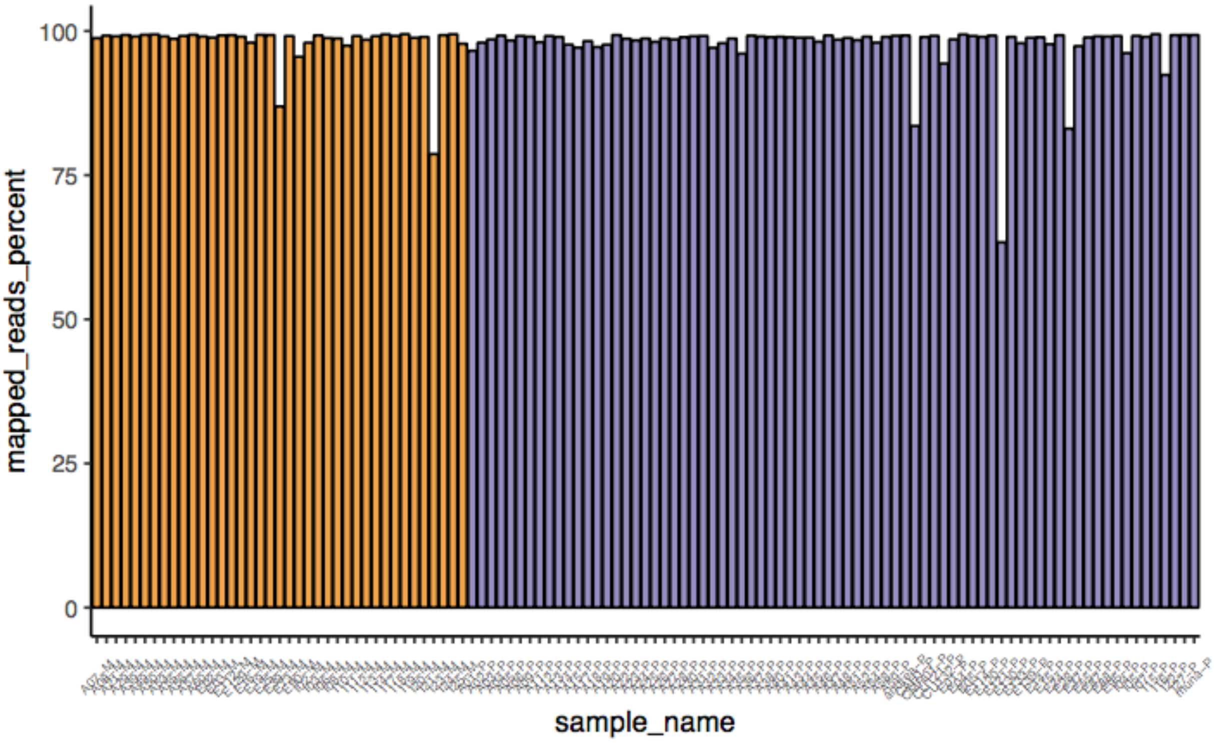

**Supplementary Figure 3: Mapped read proportion by social type.**
Proportion of reads mapping to the reference for single-queen samples in orange and multiple-queen samples in purple. There is no significant difference between the means of proportion of read mapping between social types (T-test, *P* = 0.58).

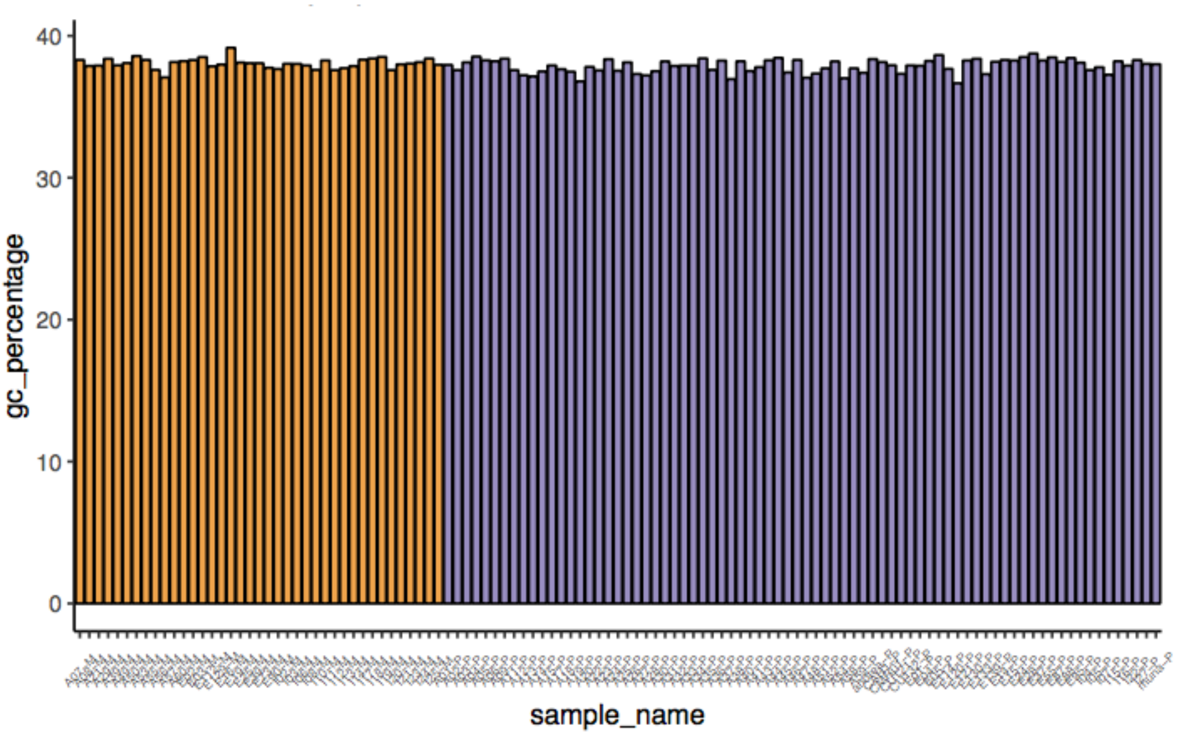

**Supplementary Figure 4: GC content proportion by social type.**
Data from Qualimap report. There is no significant difference between the means of GC content between social types (T-test, *P* = 0.023).

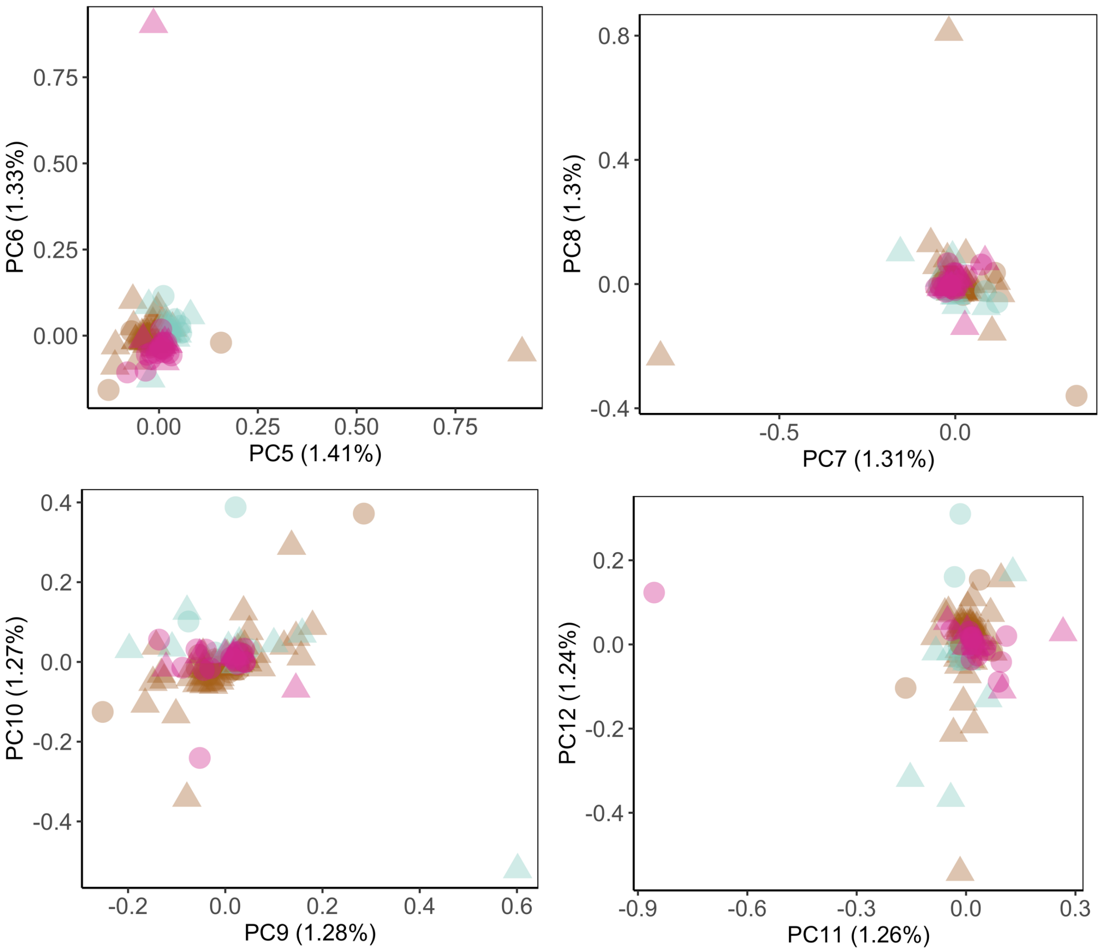

**Supplementary Figure 5: PCA for minor PCs**

Samples do not cluster by social type (triangle for multiple-queen colony, round for single-queen colony). PC analysis from variance-standardised relationship matrix of 121,786 within-population polymorphic SNPs, supported by 75% of samples; 108 samples from Bruniquel (in brown), Vigliano (in pink), Iberia (in blue).

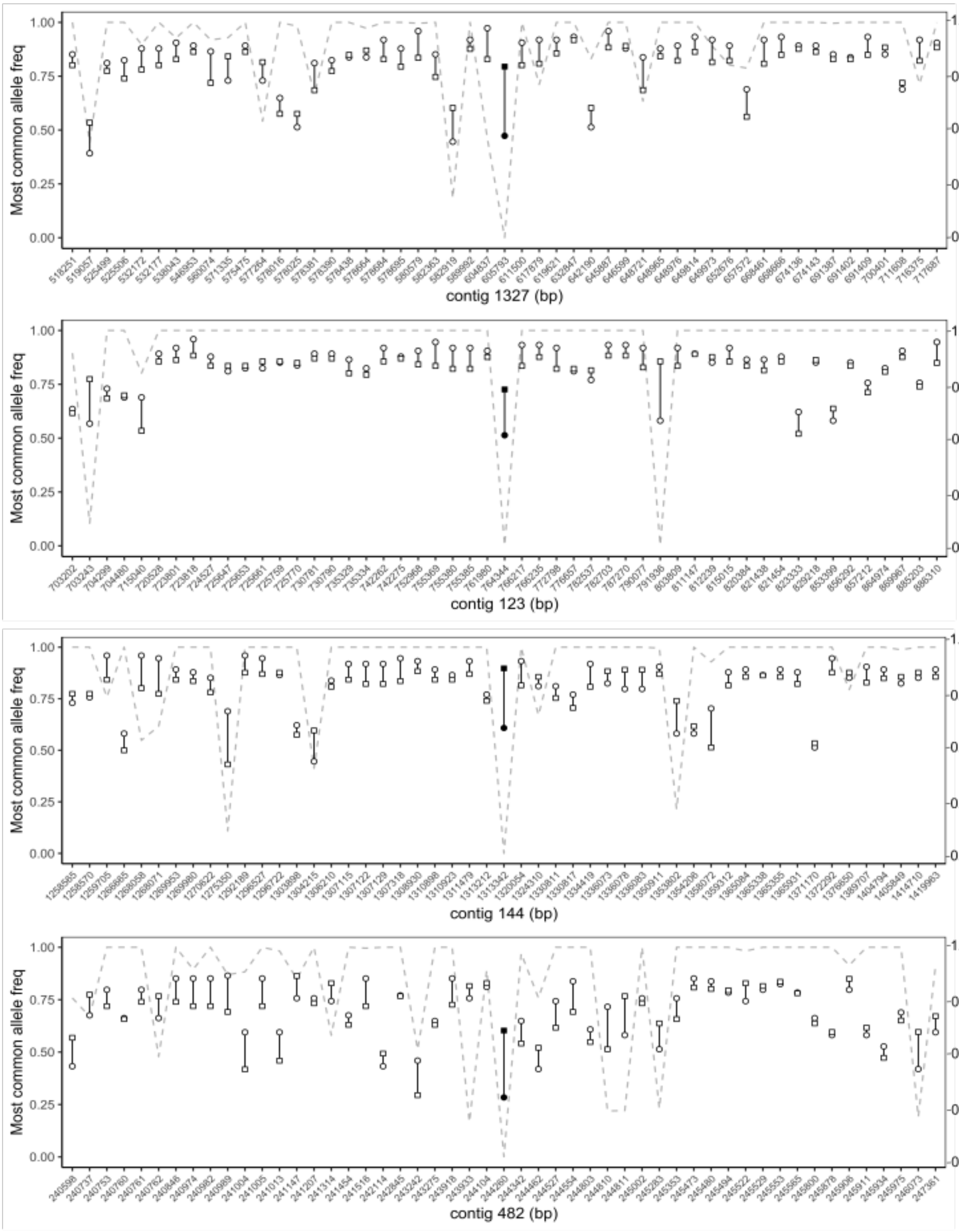

**Supplementary Figure 6: Frequency of most common allele (continued below)**

Each fragment of contig (x axis) contains the SNP significantly associated with social form (point filled in black) and up to 50 neighbouring SNPs, taken from the analysis of association with social type. For each locus, the frequency of the most common allele (y axis, left) is calculated for the single-queen samples (round point) and the multiple-queen samples (square point). The dashed line is the adjusted *P* value (y axis, right) from the association test for social type (Fisher's exact test, Bonferroni correction).

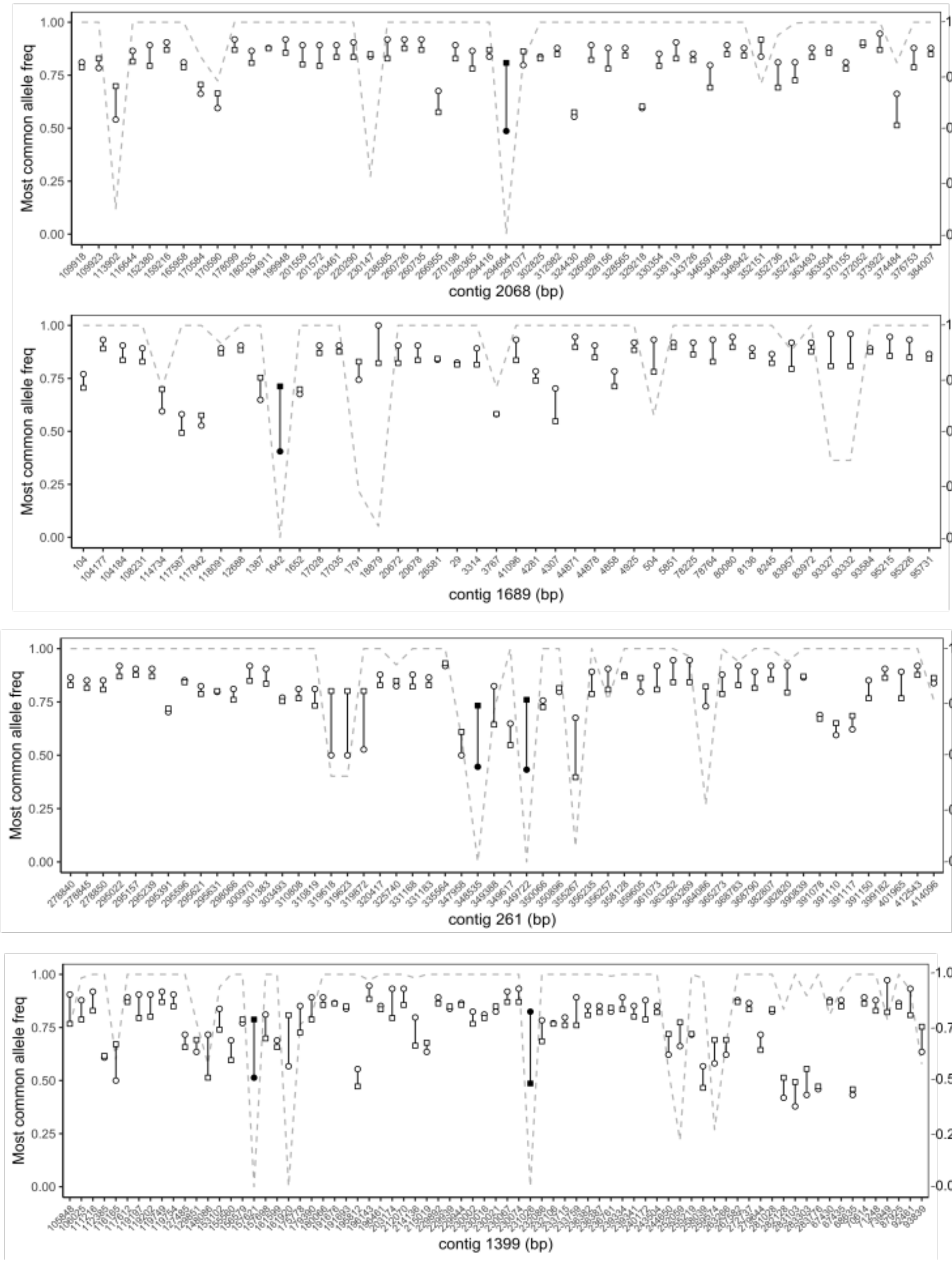

**Supplementary Figure 6 (continued): Frequency of most common allele**

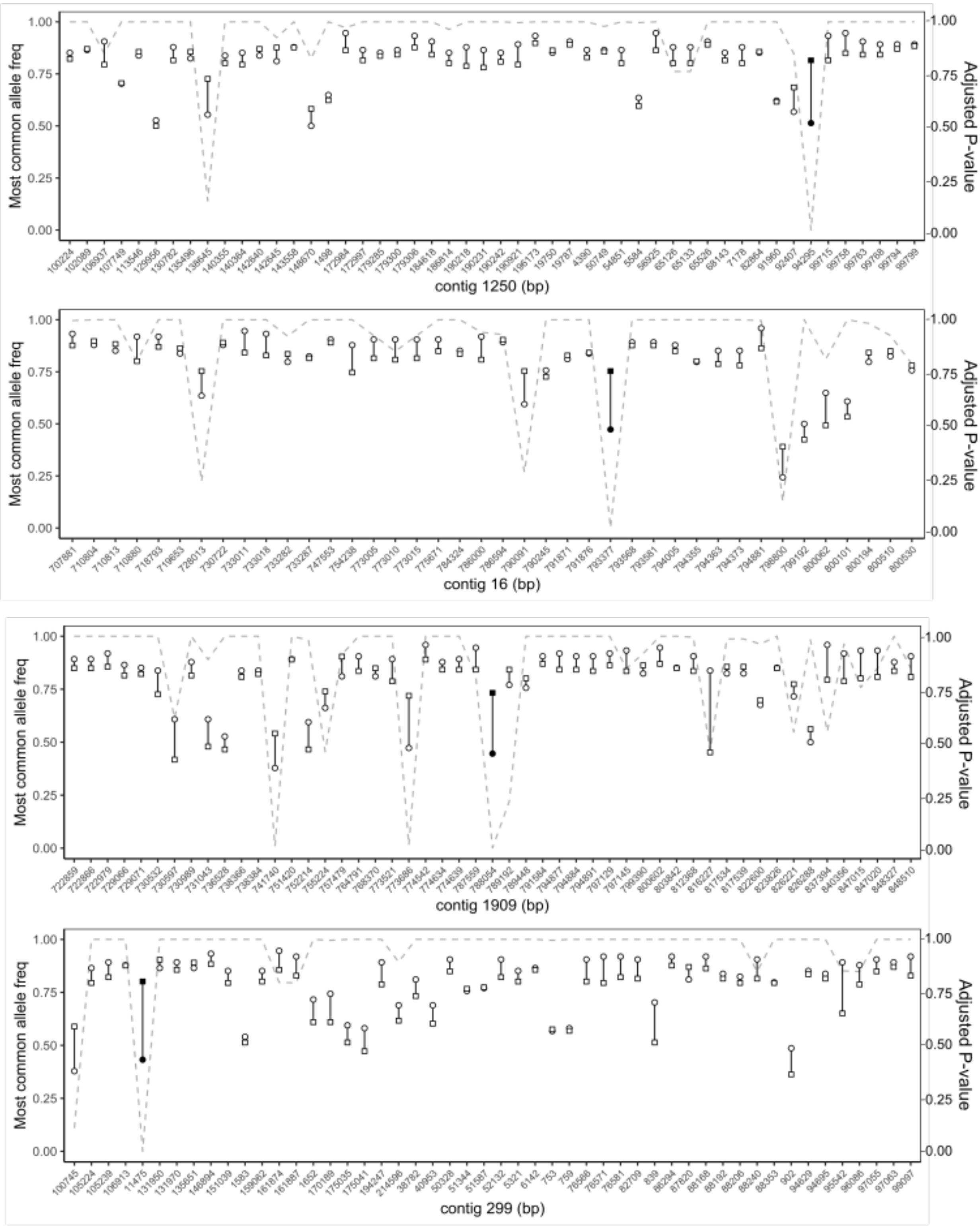

**Supplementary Figure 6 (continued): Frequency of most common allele**

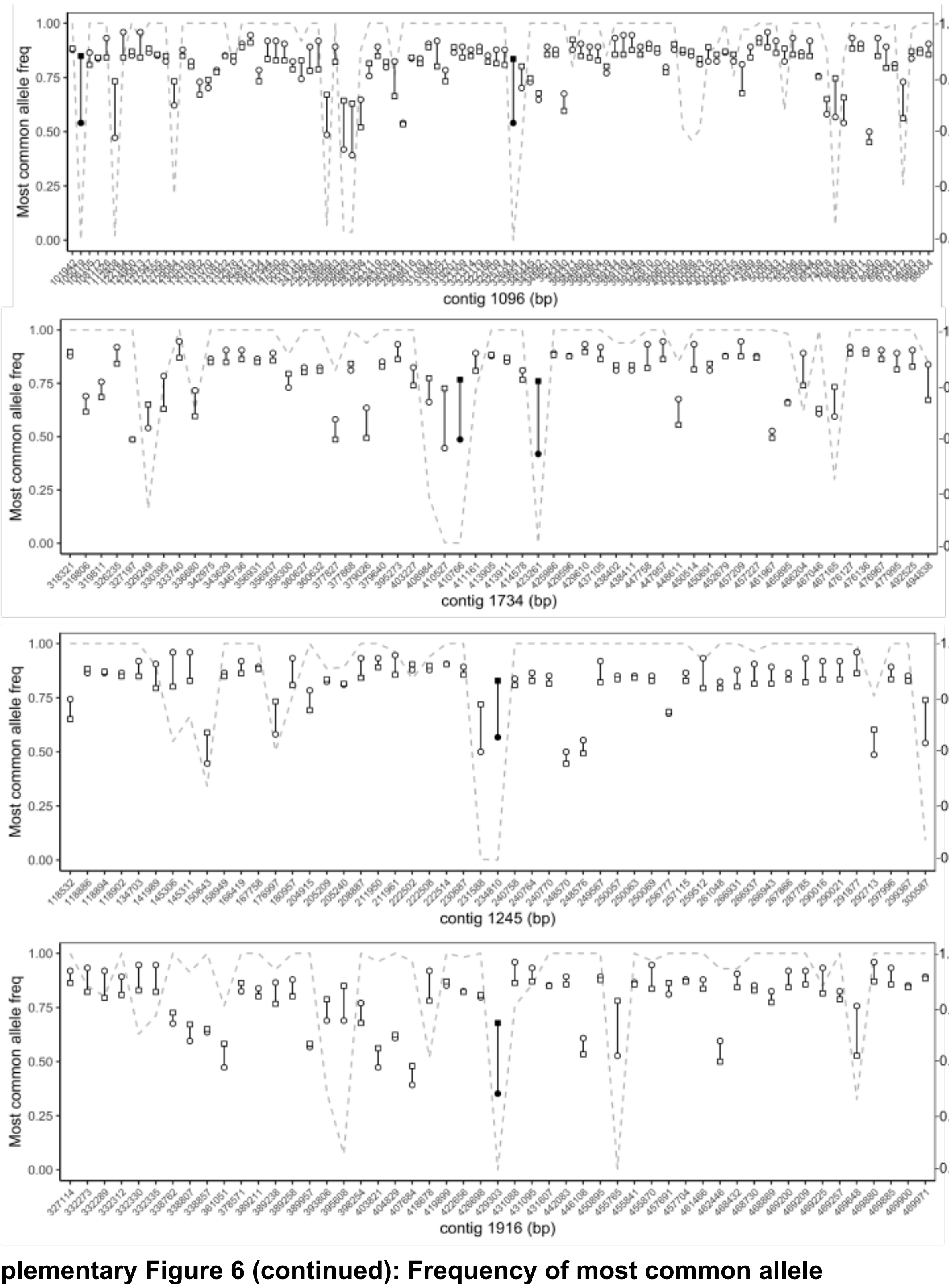

**Supplementary Figure 6 (continued): Frequency of most common allele**

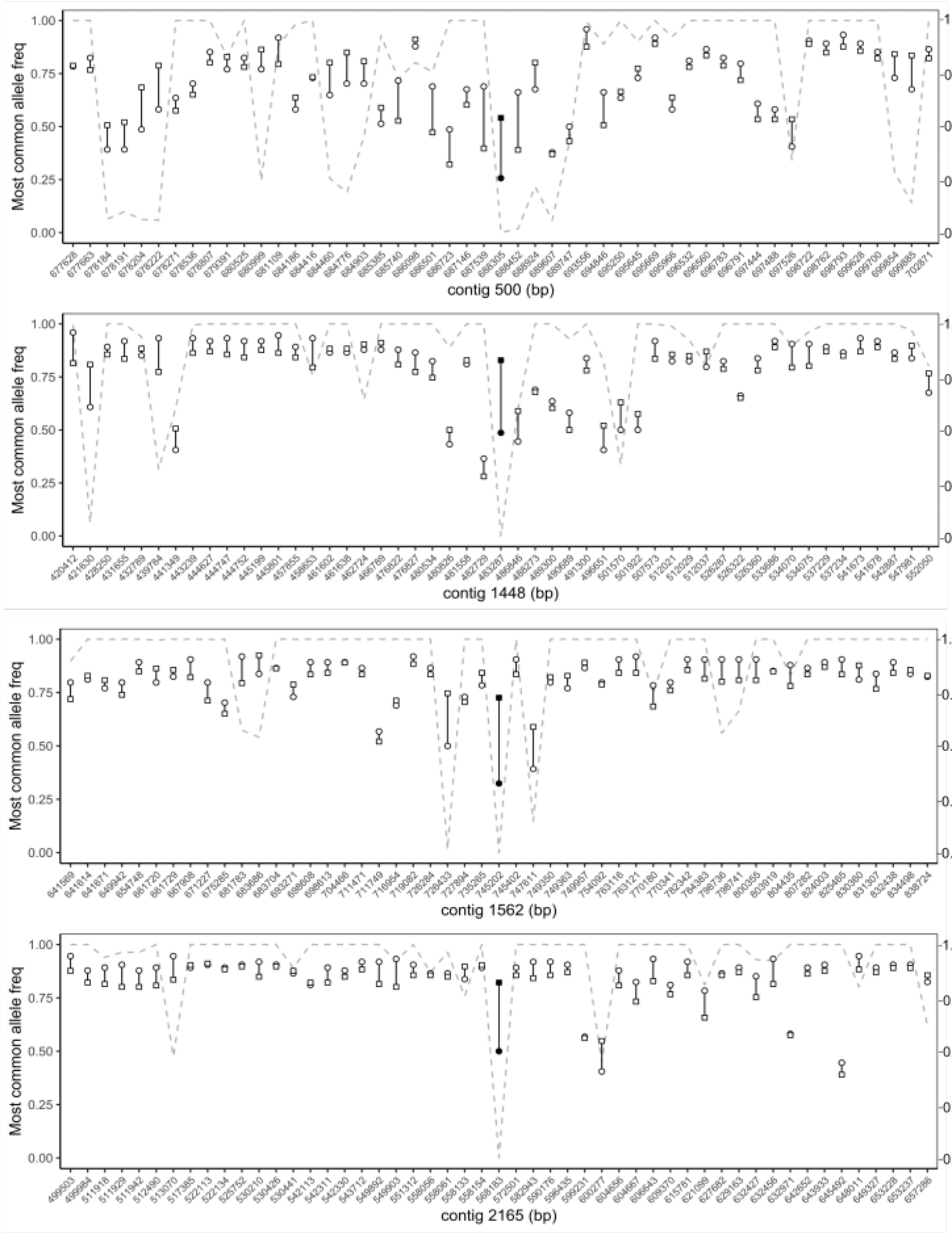

**Supplementary Figure 6 (continued): Frequency of most common allele**

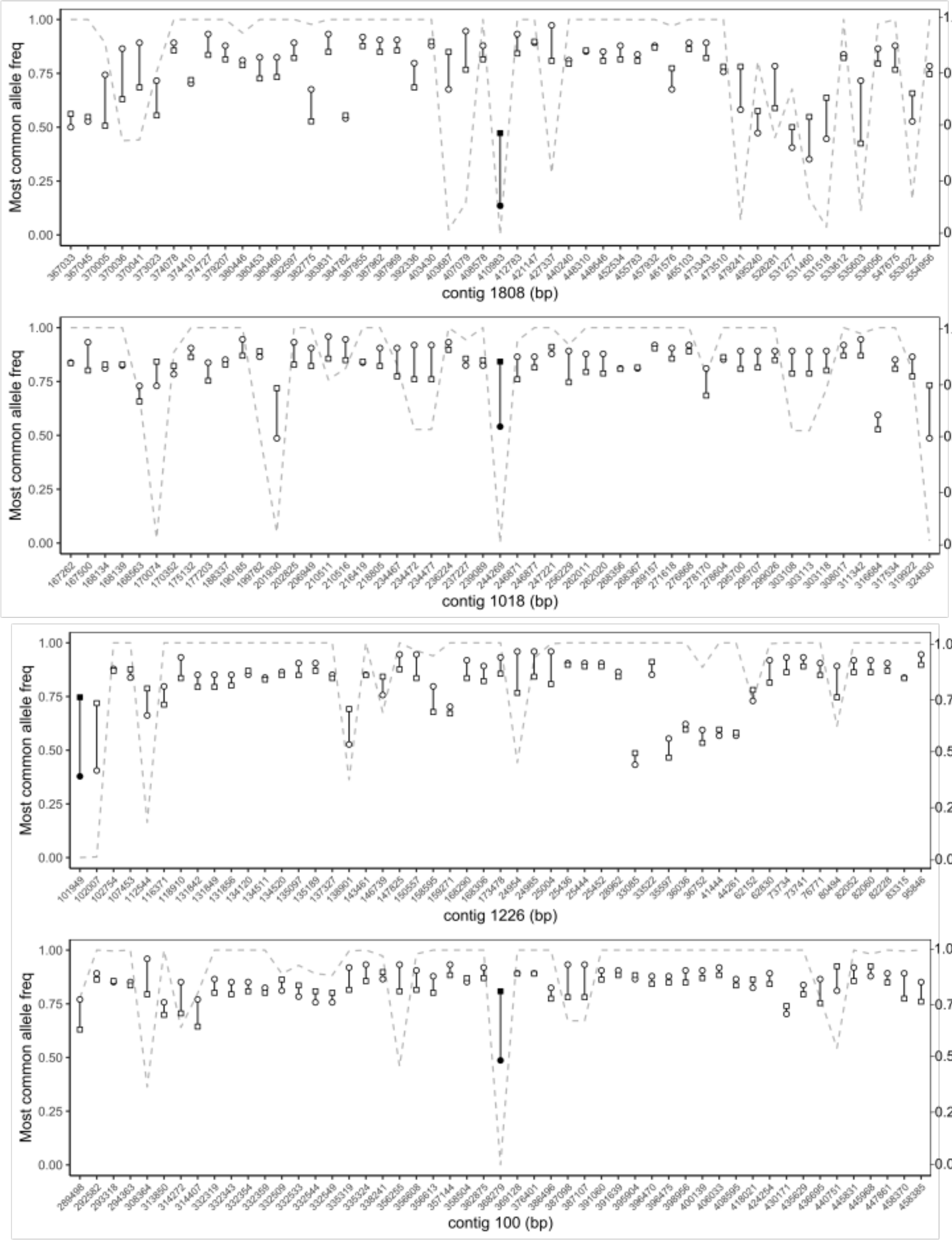

**Supplementary Figure 6 (continued): Frequency of most common allele**

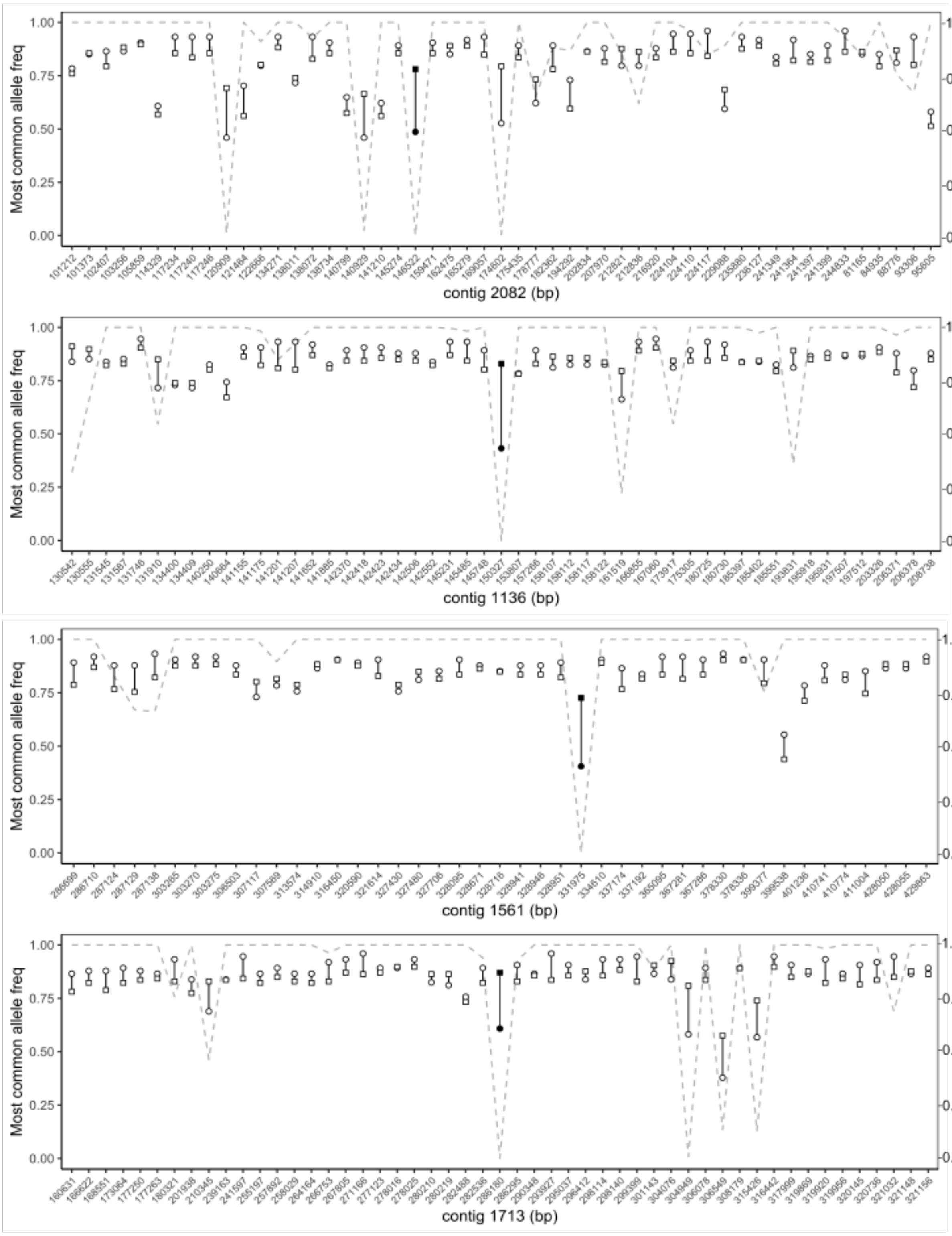

**Supplementary Figure 6 (continued): Frequency of most common allele**

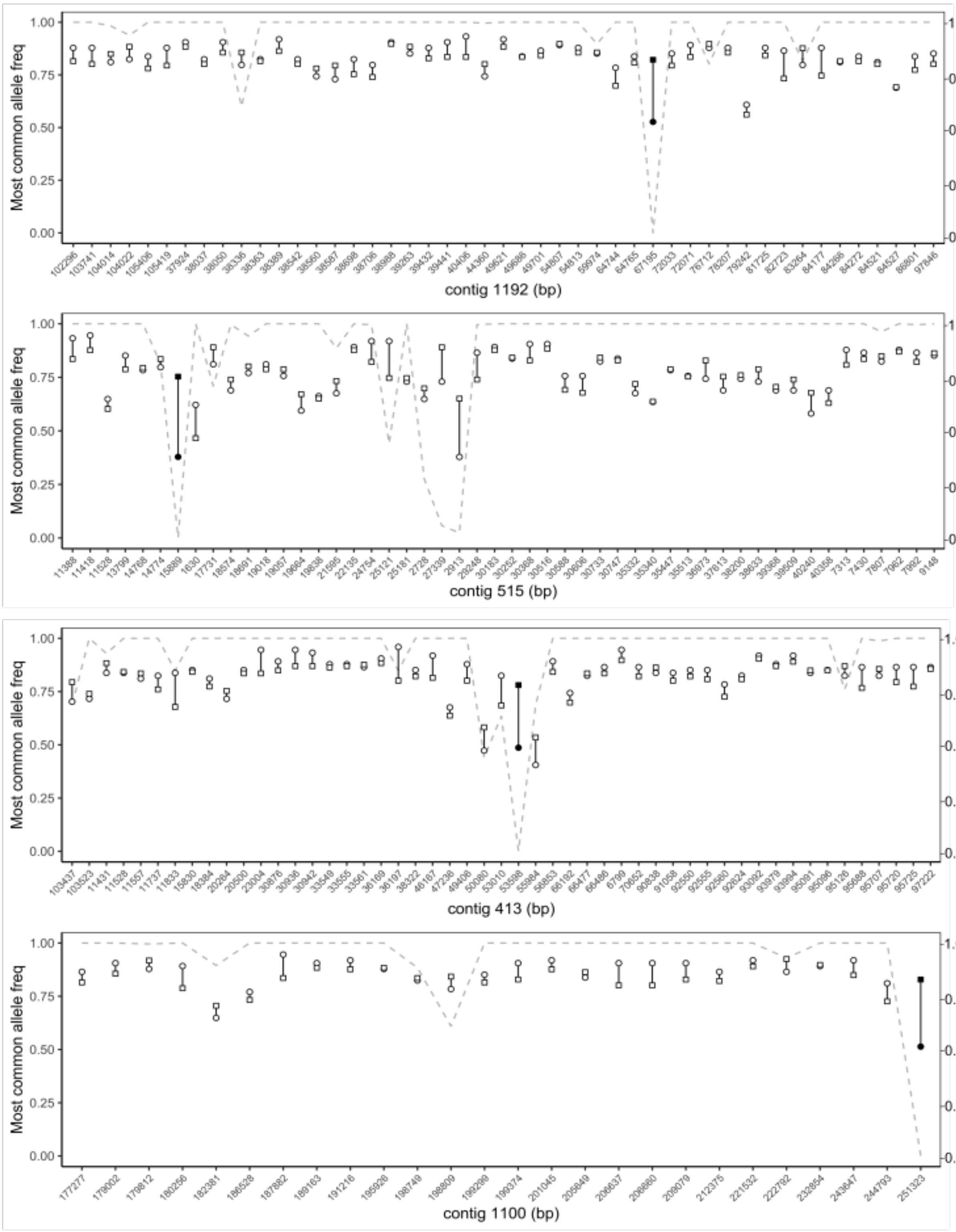

**Supplementary Figure 6 (continued): Frequency of most common allele**

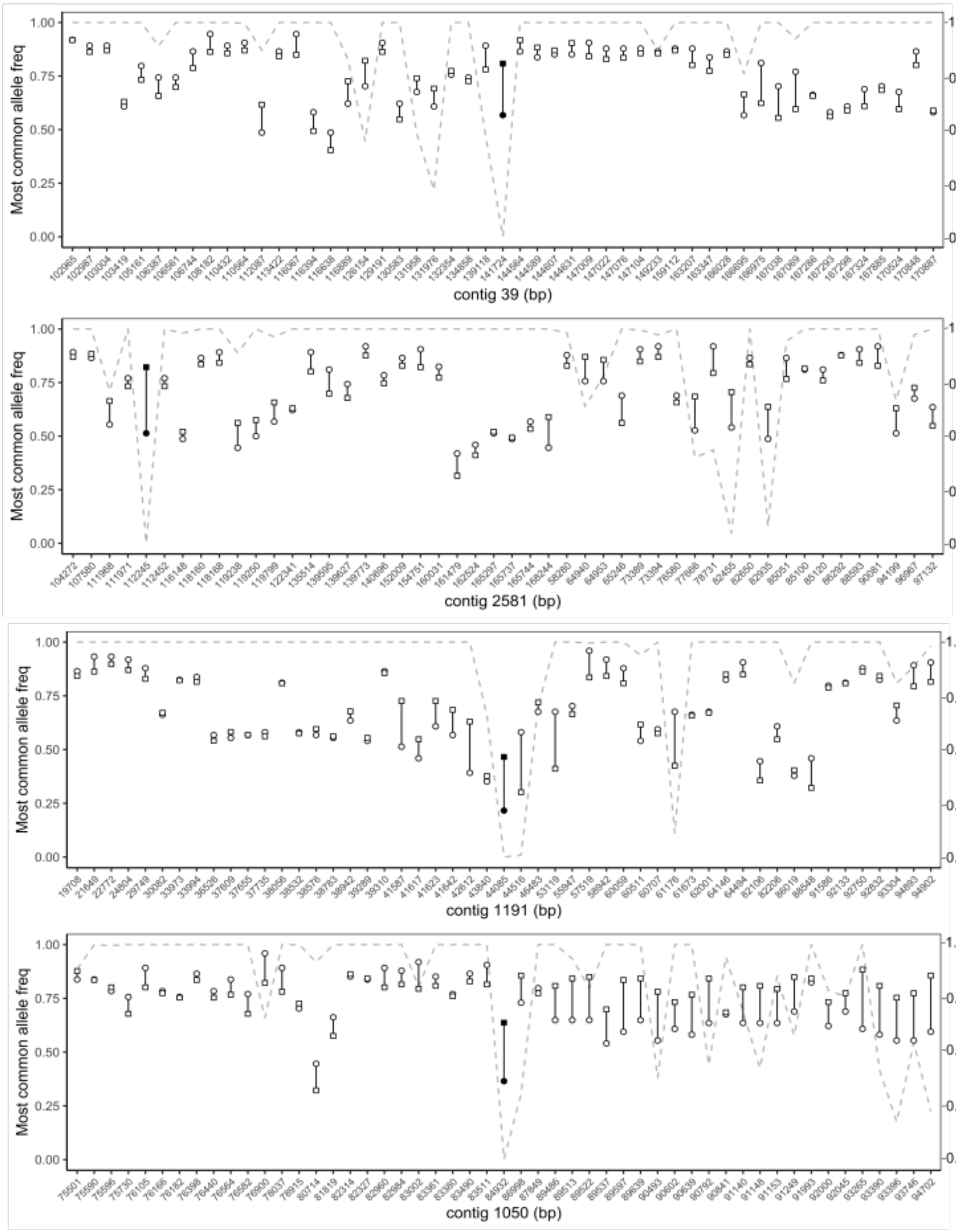

**Supplementary Figure 6 (continued): Frequency of most common allele**

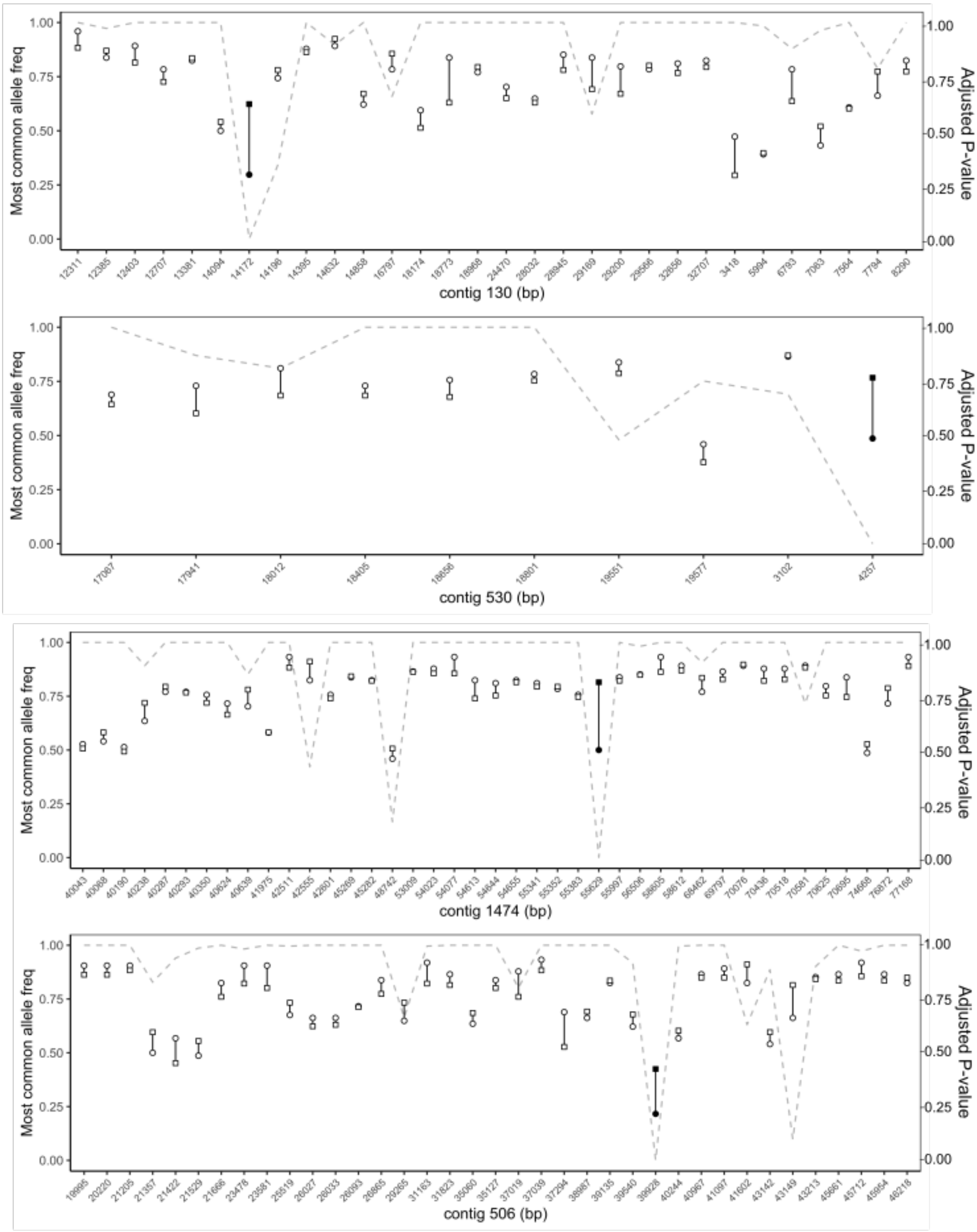

**Supplementary Figure 6 (continued): Frequency of most common allele**

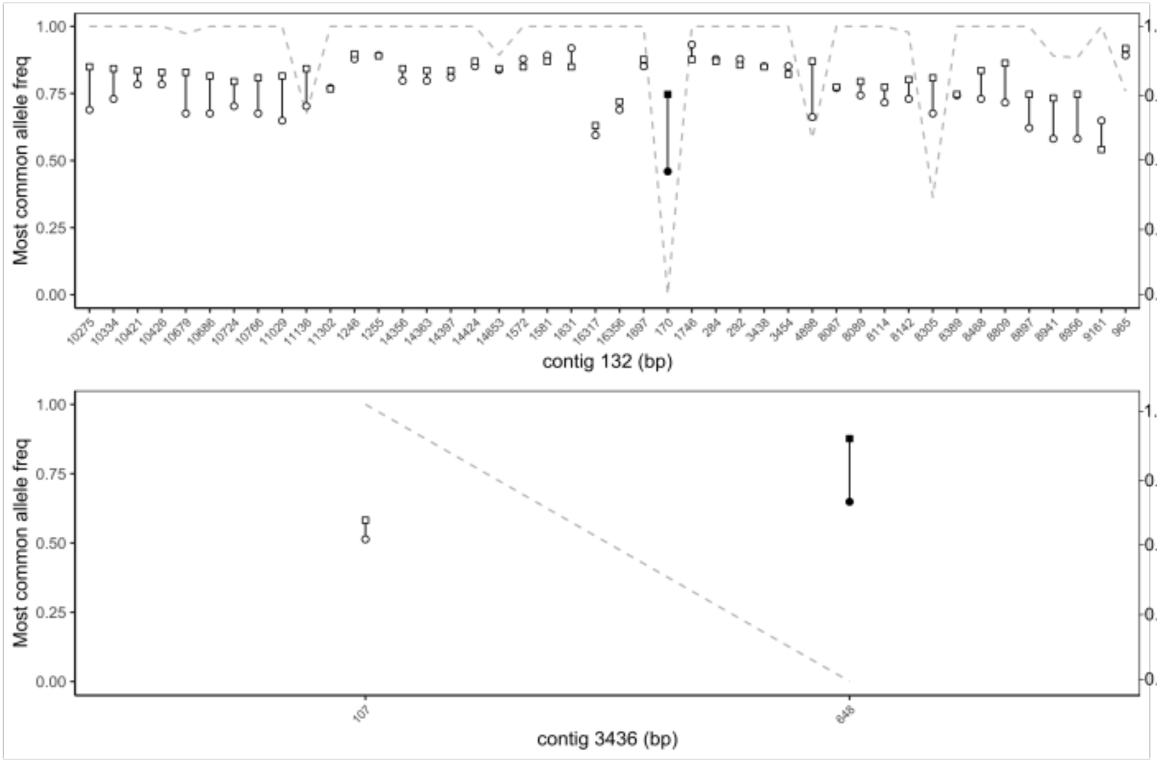

**Supplementary Figure 6 (continued): Frequency of most common allele**

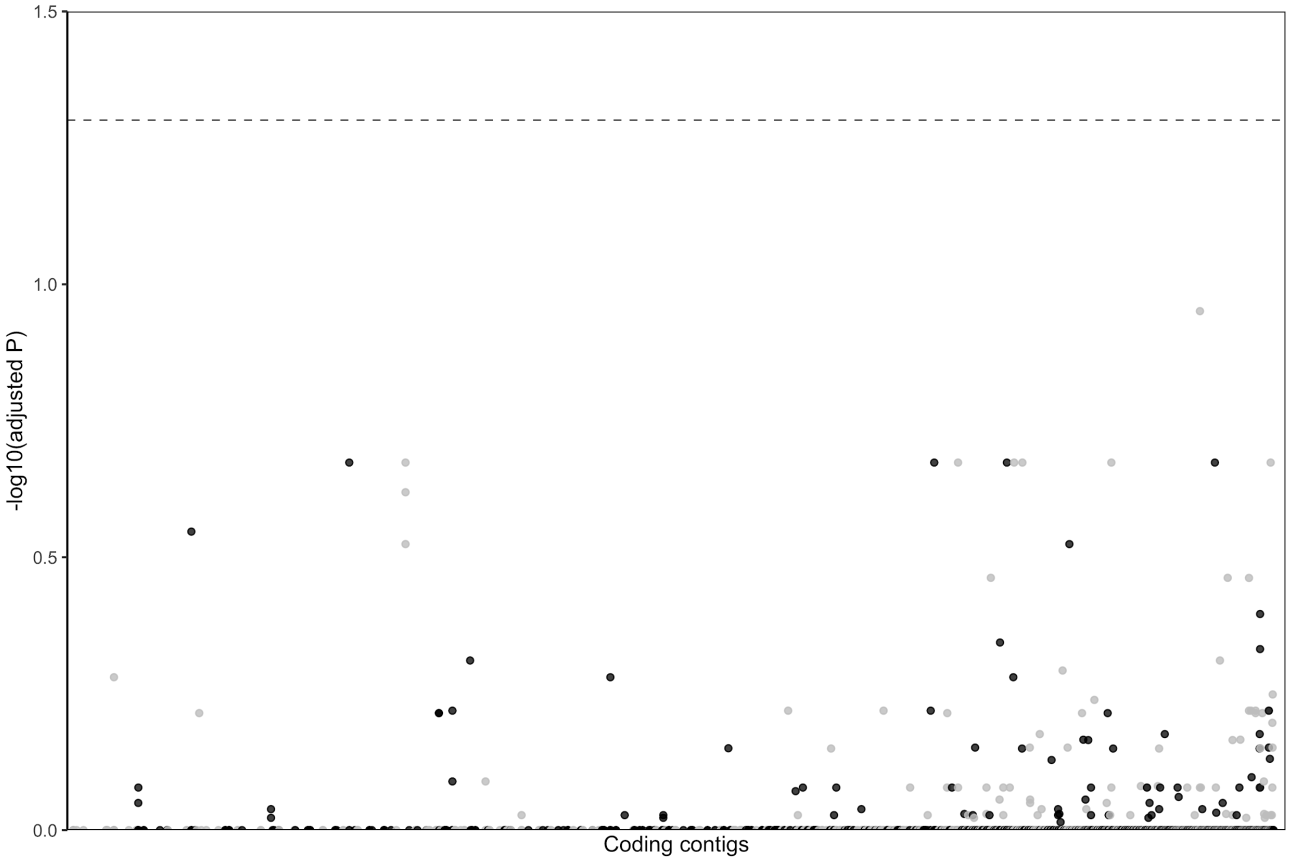

**Supplementary Figure 7: No association with social form in coding regions.**
No SNP is significantly associated with social form after multiple comparisons adjustment, among 3,360 within-population polymorphic coding SNPs found in all three populations. Contigs in the x axis are ordered by length. Horizontal dashed line represents the significance line ($P_{adj}$ = 0.05). We filtered our dataset for variants in the coding regions, using *Solenopsis invicta* protein sequences (Ensembl GFF file version 41 (Camacho et al. 2009)) as a reference in a tblastn query against Ppal_gnE assembly (-evalue 1e-3) (Camacho et al. 2009). We then filtered out the smaller fragments in the output (< 300 aa), obtaining a *Pheidole* BED file from these results. We then filtered out variants from the original VCF that are not in the newly created BED file with BEDtools v2.26.0 intersect. We obtained a SNP matrix and the sample names from this VCF with BCFtools query. After filtering for coding regions only, we retained 113, 234 SNPs (108 samples, within coding regions).

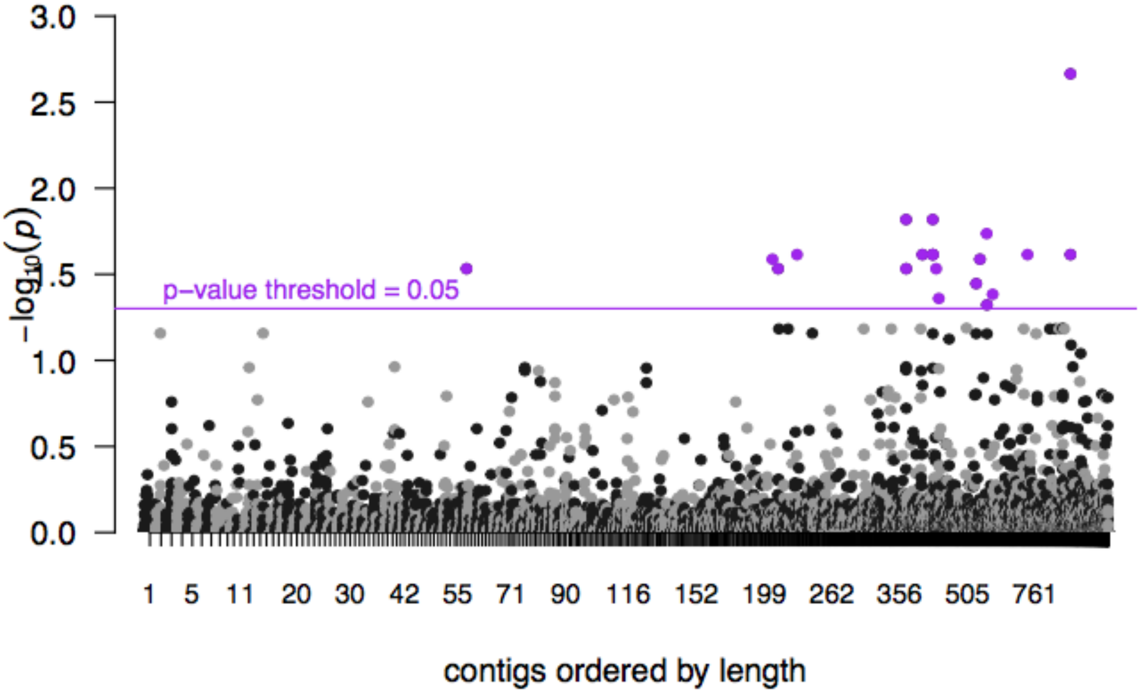

**Supplementary Figure 8: Manhattan plot for Bruniquel with 20 scattered SNPs in 16 contigs**

Association test for social type within Bruniquel population. Total number of SNPs: 97,684 SNPs. The contigs are ordered by length.

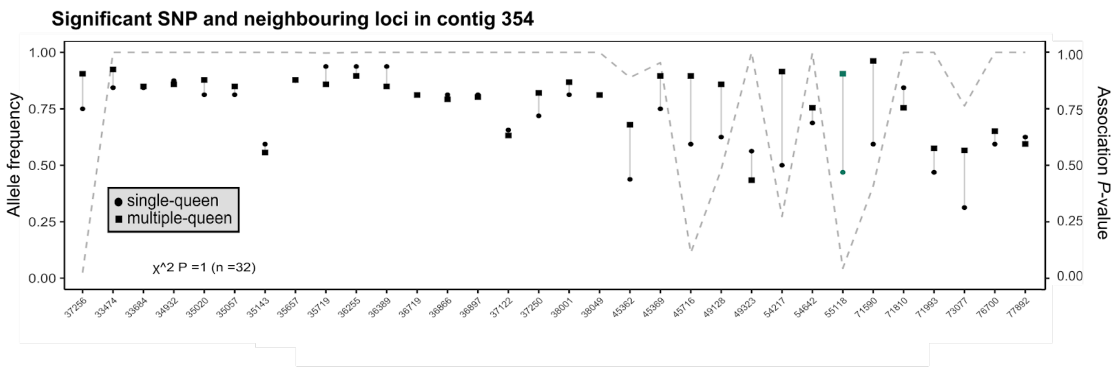

**Supplementary Figure 9: Frequency of most common allele of 20 SNPs that are significantly associated with social form and their neighbouring loci within the Bruniquel population (continued below).**
Each fragment of contig contains the SNP significantly associated with social form (turquoise) and up to 50 neighbouring SNPs in the whole dataset (coding and non-coding regions, within-Bruniquel loci). For each locus, the frequency of the most common allele is calculated for the monogynous samples (round) and the polygynous samples (square). At the contig level, we assessed the probability that each observation (frequencies of SNPs that are: non-significant monogynous, non-significant polygynous, significant monogynous, significant polygynous) falls in the corresponding class (Chi-square test)

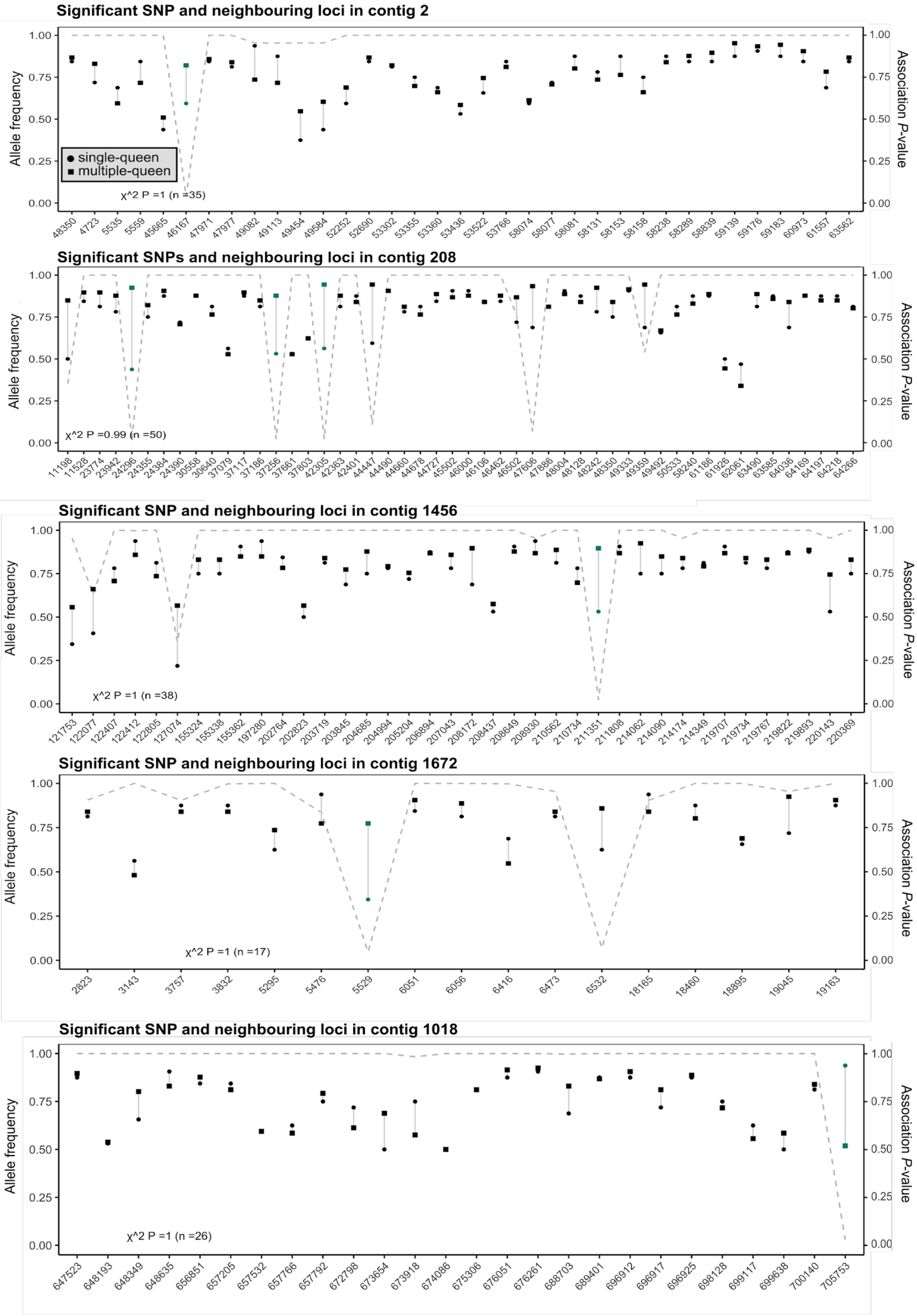

**Supplementary Figure *9* (continued): Frequency of most common allele of 20 SNPs that are significantly associated with social form and their neighbouring loci within the Bruniquel population**

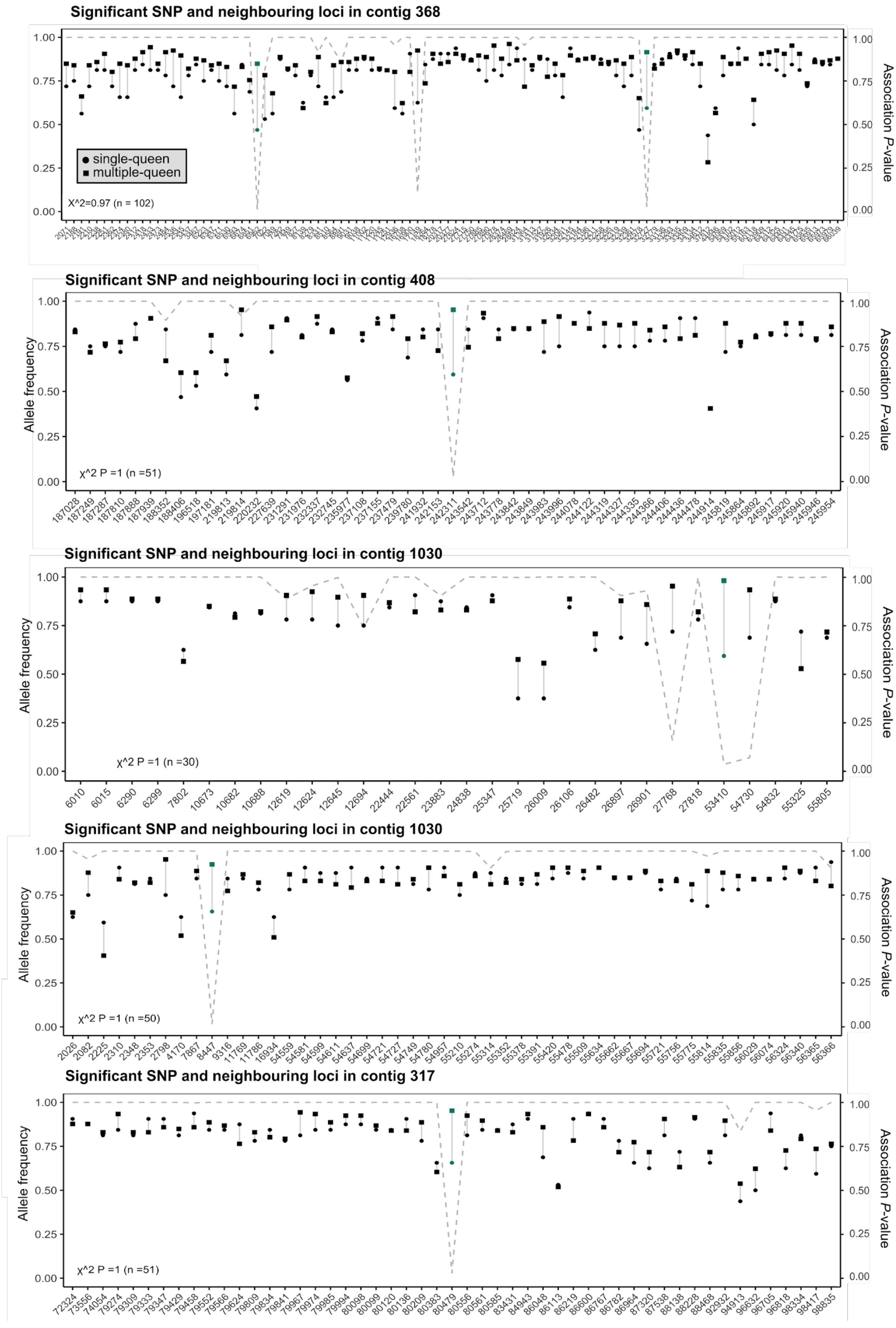

**Supplementary Figure 9 (continued): Frequency of most common allele of 20 SNPs that are significantly associated with social form and their neighbouring loci within the Bruniquel population**

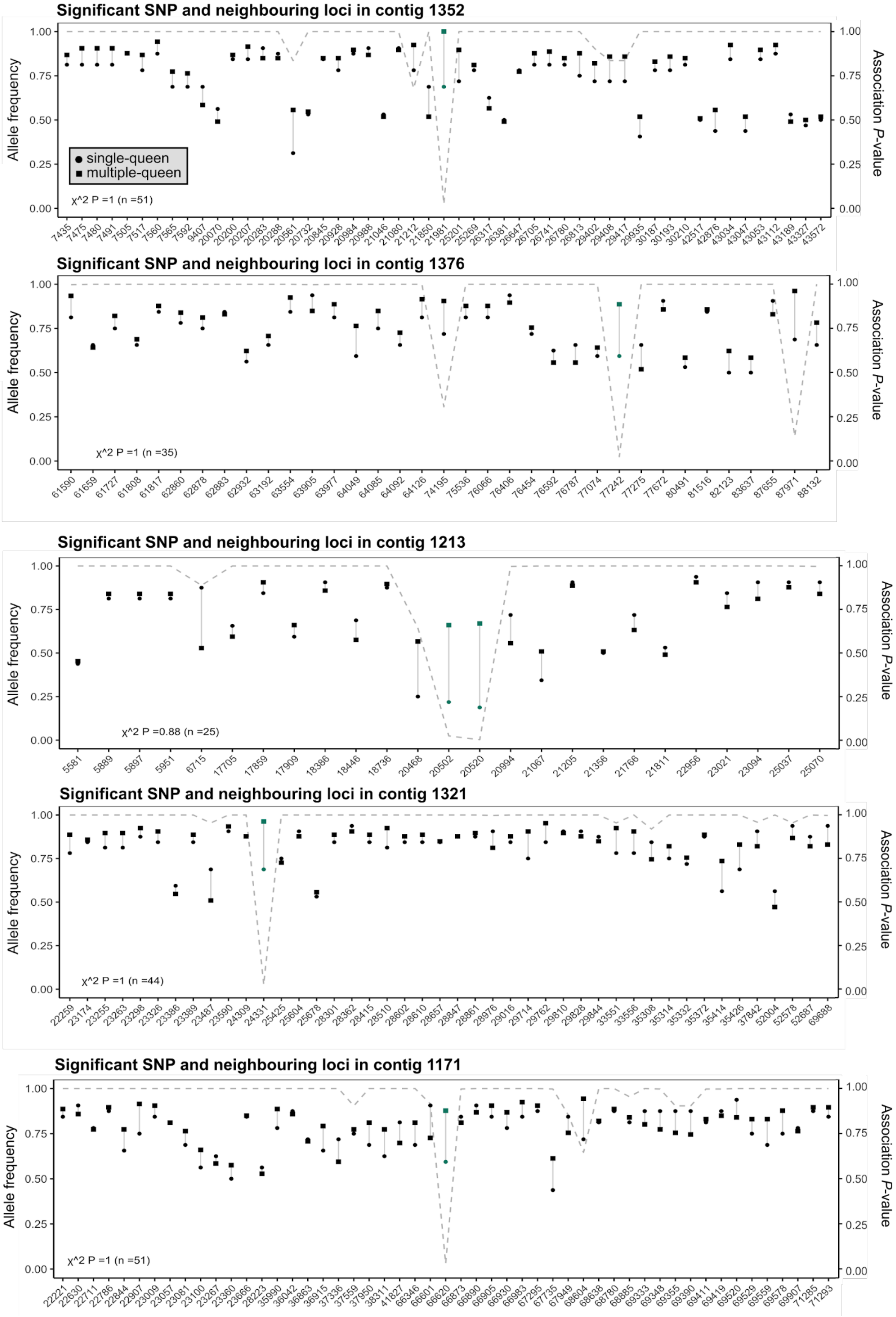

**Supplementary Figure 9 (continued): Frequency of most common allele of 20 SNPs that are significantly associated with social form and their neighbouring loci within the Bruniquel population**

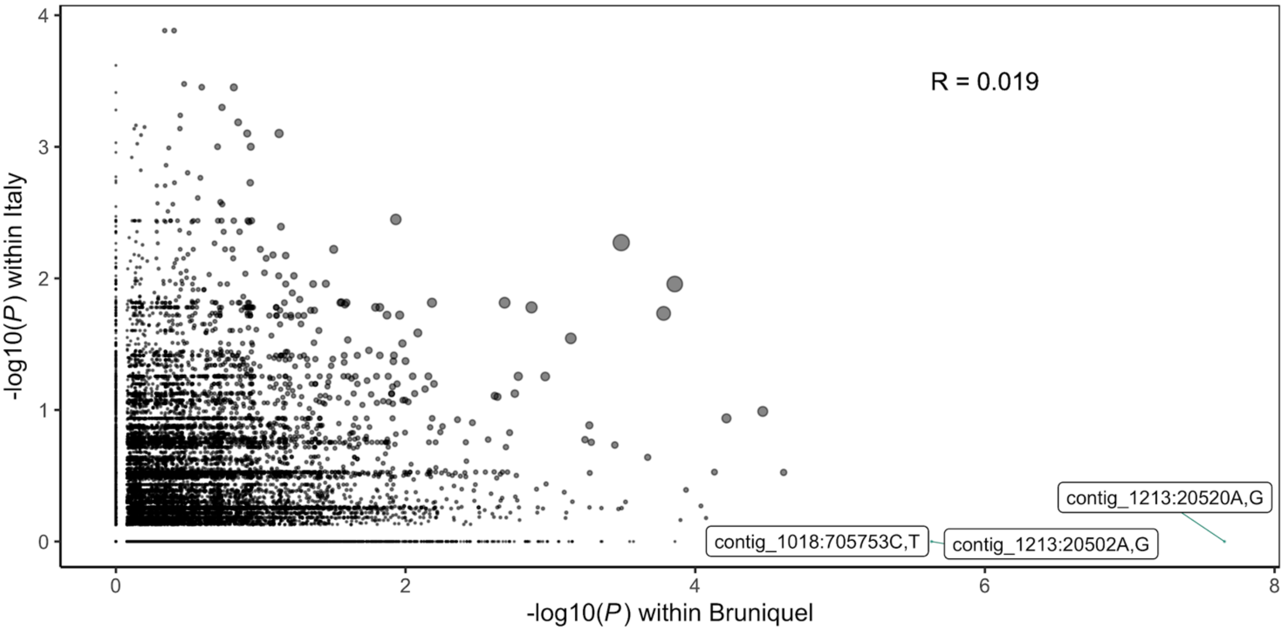

**Supplementary Figure 10: Weak correlation between association with social organisation between Bruniquel and Vigliano population**

Weak correlation between association with social organisation between Bruniquel and Vigliano population (Pearson's Correlation on unadjusted *P* values, r = 0.44, t = 20.651, df = 1746, p-value < 2.2e-16). The x axis is the raw *P* value (-log10) for Bruniquel SNPs. The y axis is the raw *P* value (-log10) for Vigliano SNPs. The points represent loci. The circle radius size equals the multiplication of absolute raw *P* values. Only three loci are significantly associated with social organisation in Bruniquel population (points with labels related to location and allele variation).

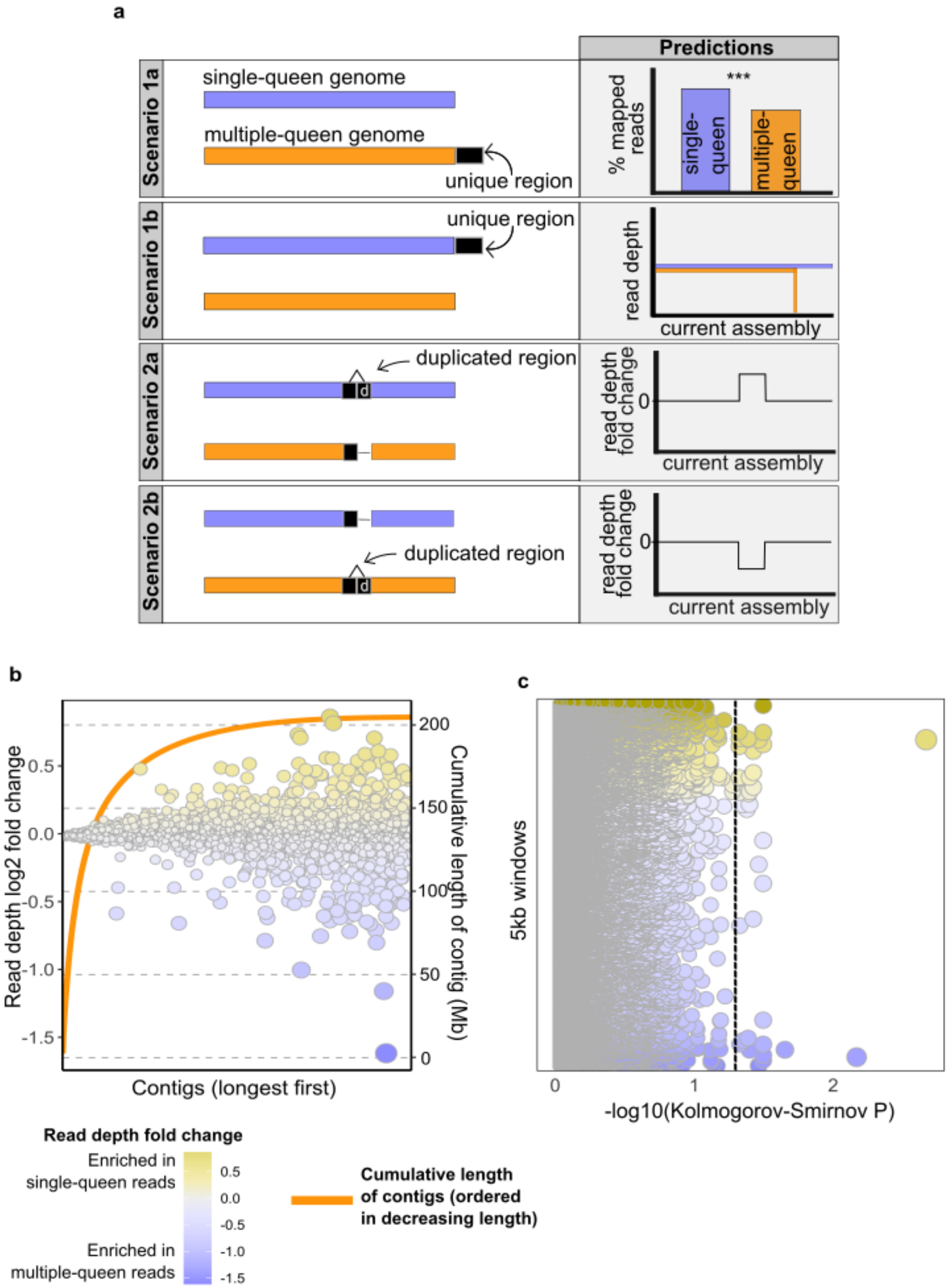

**Supplementary Figure 11: No coverage skew between social forms (contigs and 5kb regions)**

**a)** Scenarios for potential reference skew. **Scenario 1a**: the multiple-queen genome contains a unique region that does not map to the single-queen assembly, thus the proportion of mapped reads over the total number of reads is lower in multiple-queen samples than in single-queen samples. **Scenario 1b**: the unique region is in the single-queen genome, thus no multiple-queen reads will map to this region. **Scenario 2a**: the

duplicated region is in the single-queen genome, thus there are more single-queen reads mapping this region (positive read depth fold change single-queen over multiple-queen).
**Scenario 2b**: the duplicated region is in the multiple-queen genome, thus there are more multiple-queen reads mapping this region (negative read depth fold change).
**b**) The majority of the assembly (size represented in orange line) is made of contigs with an equal proportion of reads from single- and multiple-queen colonies (contig read depth represented in points for the contig-level fold change of coverage in multiple-queen samples with respect to single-queen samples). Some contigs are enriched in mapped reads from single-queen samples (top, in yellow), other contigs are enriched in mapped reads from multiple-queen samples (bottom, in purple). Any point (= one contig) with a $\log_2$ fold change of 0 (in white) represents the genome-wide average of fold change coverage of each contig by mapped reads. The circle radius size equals the deviation from the genome-wide average fold change.
**c)** 5-kb-level fold change of coverage in multiple-queen samples with respect to single-queen samples. Each contig is represented fragmented in 5kb windows (one point = one window). Each 5kb window has been investigated for significant difference in read coverage between social forms (Kolmogorov-Smirnov test, $P_{adj} = 0.05$). The dotted line denotes the significance level that separates 5kb regions with significant difference in read coverage between social forms (right side). The colour gradient describes the enrichment: purple windows from contigs that are enriched in multiple-queen reads (bottom), yellow windows from contigs are enriched in single-queen reads (top).

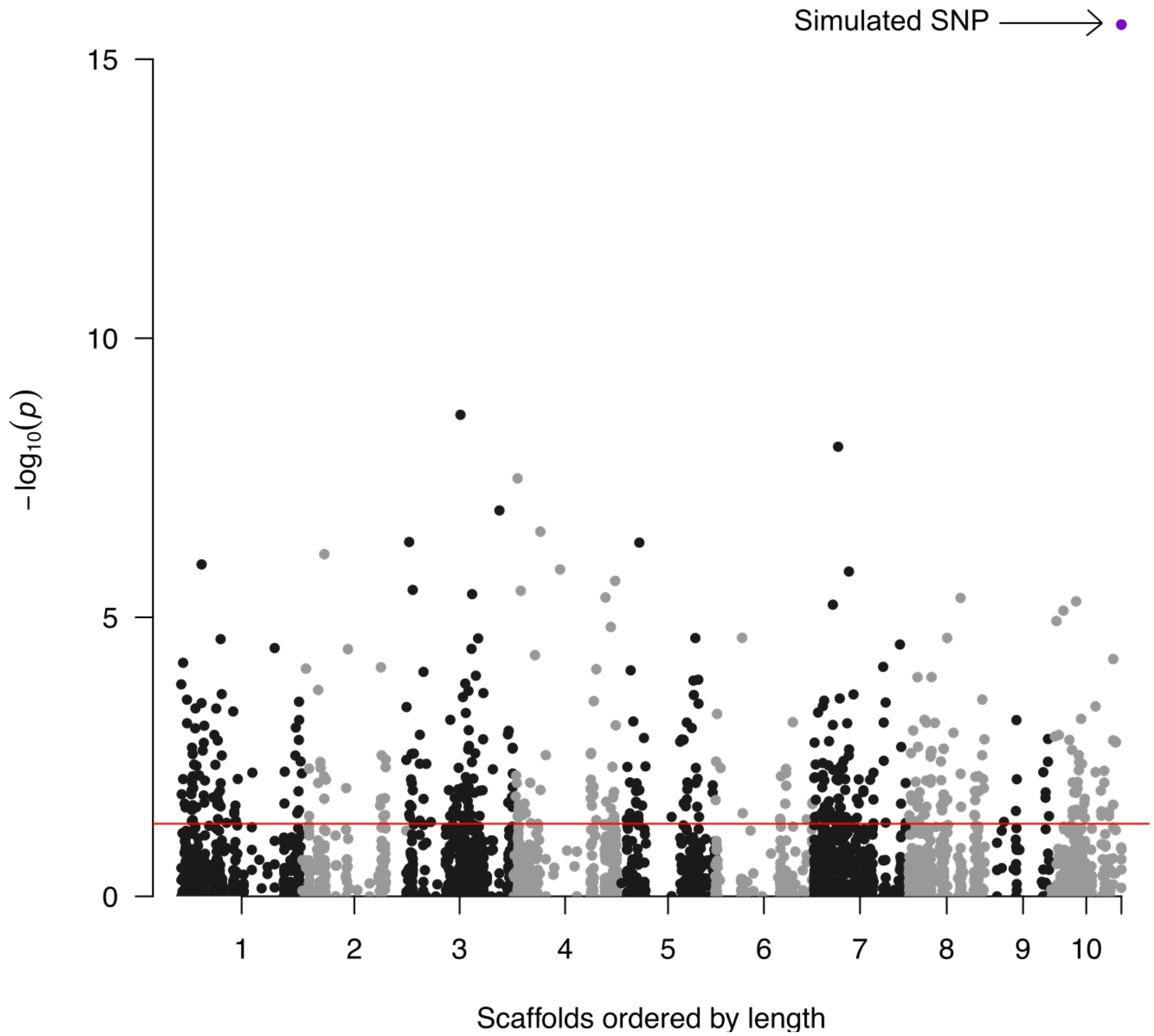

**Supplementary Figure 12: Purple simulated SNP at the top right is the most significant variant in Fisher's exact tests.**

Monogynous samples are homozygous at this locus, polygynous samples are heterozygous. Red horizontal line shows the *P* value threshold (0.05). All other SNPs are from real data (Bonferroni correction).

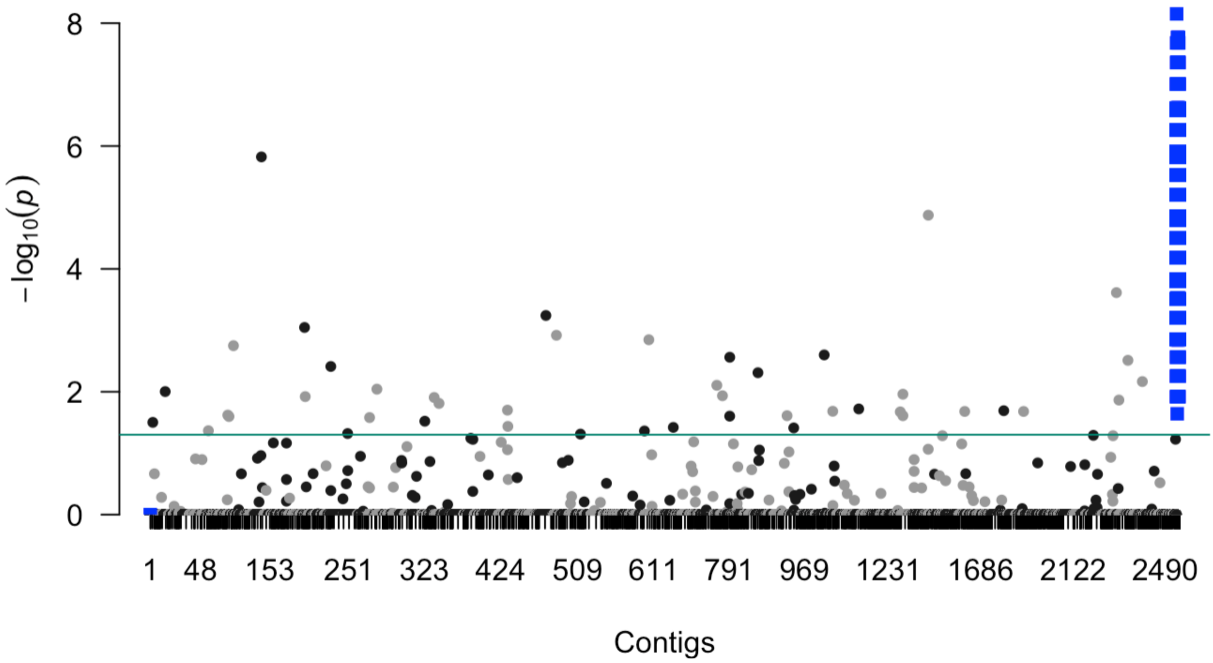

**Supplementary Figure 13: *Solenopsis invicta* simulation of supergene detection with Fisher's exact test of allele count.**

124, 840 association tests for social type (Bonferroni correction) for 121,786 real SNPs and 3,054 simulated SNPs. The 121,786 real *Pheidole* SNPs are supported by 75% samples, within-population polymorphic (see main text analysis). The 3,054 simulated SNPs (blue squares) reflect the *Solenopsis* system: all monogynous samples are homozygous for the reference at the 3,054 simulated loci. A third of the polygynous samples are homozygous for the reference and two-thirds are heterozygous.

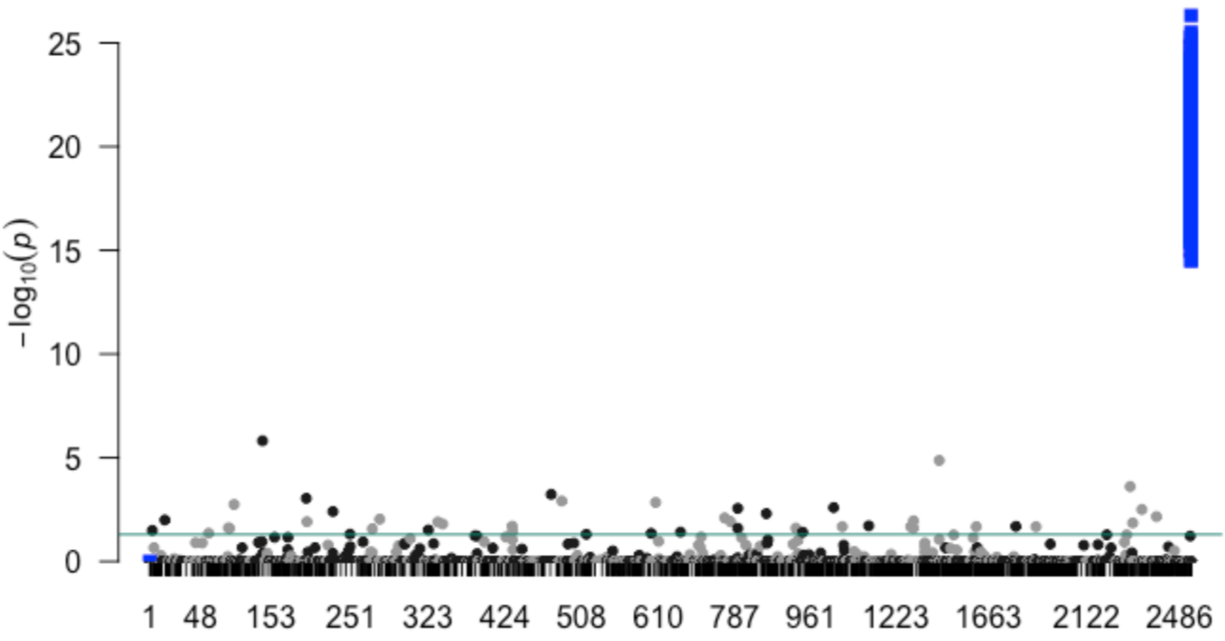

**Supplementary Figure 14: *Formica selysi* simulation of supergene detection with Fisher's exact test of allele count.**

126,291 association tests for social type (Bonferroni correction) for 121,786 real SNPs and 4,505 simulated SNPs. The 121,786 real *Pheidole* SNPs are supported by 75% samples, within-population polymorphic (see main text analysis). The 4,505 simulated SNPs (blue squares) reflect *Formica* system: all monogynous samples are homozygous for the reference at the 4,505 simulated loci. A third of the polygynous samples are homozygous for the alternative and two-thirds are heterozygous.

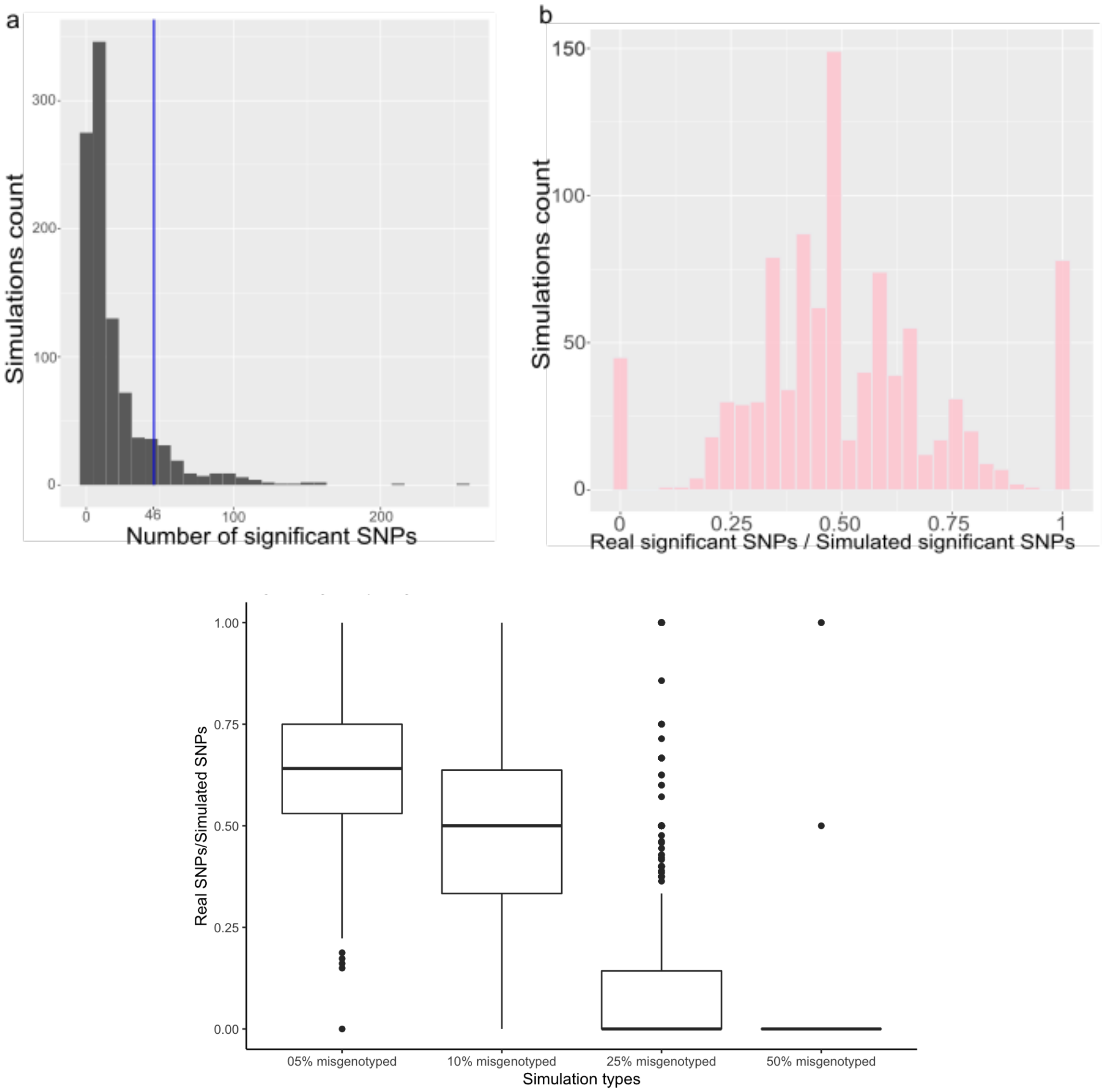

**Supplementary Figure 15: Misgenotyping simulations**

a) Histogram of the numbers of SNPs that are significantly associated with social form, over 1000 simulations, in which 10% of the samples are misgenotyped (i.e., assigned the alternative social type). Significance is measured by Fisher's exact tests and Bonferroni correction. The 48 real SNPs are indicated with the blue line. Most simulations contain less SNPs with significant association than the real dataset (regardless of the exact loci).
b) The association analysis is powerful enough to include 10% of mislabelling the colonies' social type. Indeed, for more than 950 simulations, the real SNPs with significant association (n = 48) are recovered in the simulated SNPs with social association set.
c) Proportion of real SNPs, with social association, recovered by misgenotyping simulations for 5% to 50% of samples being assigned the wrong labels. Assuming that our analysis contains up to 10% of misgenotyping, the analysis design (GWAS using Fisher's exact test and Bonferroni correction) is expected to recover at least 50% of true positive significant associations.
